# Identification of chaperone-independent outer membrane proteins and MtrA-assisted MtrB folding in *Shewanella oneidensis*

**DOI:** 10.1101/2025.06.06.657845

**Authors:** Lukas Kneuer, Bente Siebels, Lucas Marin, Laura-Alina Philipp, Johannes Gescher

## Abstract

Extracellular electron transfer is a respiratory process conducted by a number of microorganisms in order to access insoluble or membrane impermeable electron acceptors or donors. The process has wide implication for the biogeochemistry of our planet and offers many opportunities for biotechnological applications. Outer membrane spanning electron transfer is conducted by the model organism *Shewanella oneidensis* MR-1 by a stable trimeric protein complex. The electron conduit consists of two *c*-type cytochromes on either site of the outer membrane and a β-barrel protein in the middle that seems to facilitate interaction of the two heme containing proteins. This study reveals that the periplasmic *c-*type cytochrome MtrA is not only part of the electron conduit, but also assists in the periplasmic transport of the unfolded outer membrane protein MtrB, a function that was so far believed to be conducted for all outer membrane β-barrel proteins by one of the canonical chaperones SurA, Skp or DegP. However, three more ß-barrel proteins were identified that are independent of the canonical chaperones as well but still rely on the BAM complex in the outer membrane, suggesting that many more solutions for periplasmic transfer of ß-barrel protein towards the outer membrane exist in Gram-negative microorganisms.

## Introduction

A wide variety of cellular functions in organisms ranging from eukaryotes to bacteria are catalyzed by outer membrane β-barrel proteins (OMPs)^1–4^. The biogenesis of these proteins presents a fundamental biological challenge: the transport of hydrophobic structures through hydrophilic environments prior to their correct integration and folding into membrane bilayers. For decades, the prevailing paradigm has been that chaperone proteins are universally required to guide these unfolded outer membrane proteins (uOMPs) through the aqueous periplasm to prevent their premature aggregation and subsequent degradation^4–6^. In *Escherichia coli*, this process has been extensively characterized, with three periplasmic chaperones - SurA, Skp and DegP - constituting the canonical machinery responsible for uOMP trafficking^7–10^ (Fig. 1). Notably, DegP has a dual function as protease and chaperone and it is still under debate what factor would trigger the respective activity^8,11^. While these chaperones exhibit partial functional redundancy, with none being individually essential, double mutations result in synthetic or even synthetic lethal phenotypes^5^. This redundancy has been interpreted as an evolutionary safeguard, ensuring that the critical process of OMP biogenesis remains robust against the failure of individual components. Upon reaching the outer membrane, chaperone-assisted uOMPs interact with the BAM complex, which facilitates their folding and membrane insertion^6^. Current models position SurA as the primary chaperone in *E. coli*, with Skp and DegP forming a secondary pathway that rescues uOMPs that disengage from the SurA pathway^5^. However, recent studies also suggest that SurA and Skp could act synergistically to rescue aggregated uOMPs^12^. Moreover, BepA and YcaL - two ancillary proteases localized to the outer membrane - degrade β-barrel proteins that are in contact with the BAM complex in the outer membrane, but cannot be correctly integrated^13,14^.

**Figure 1.**
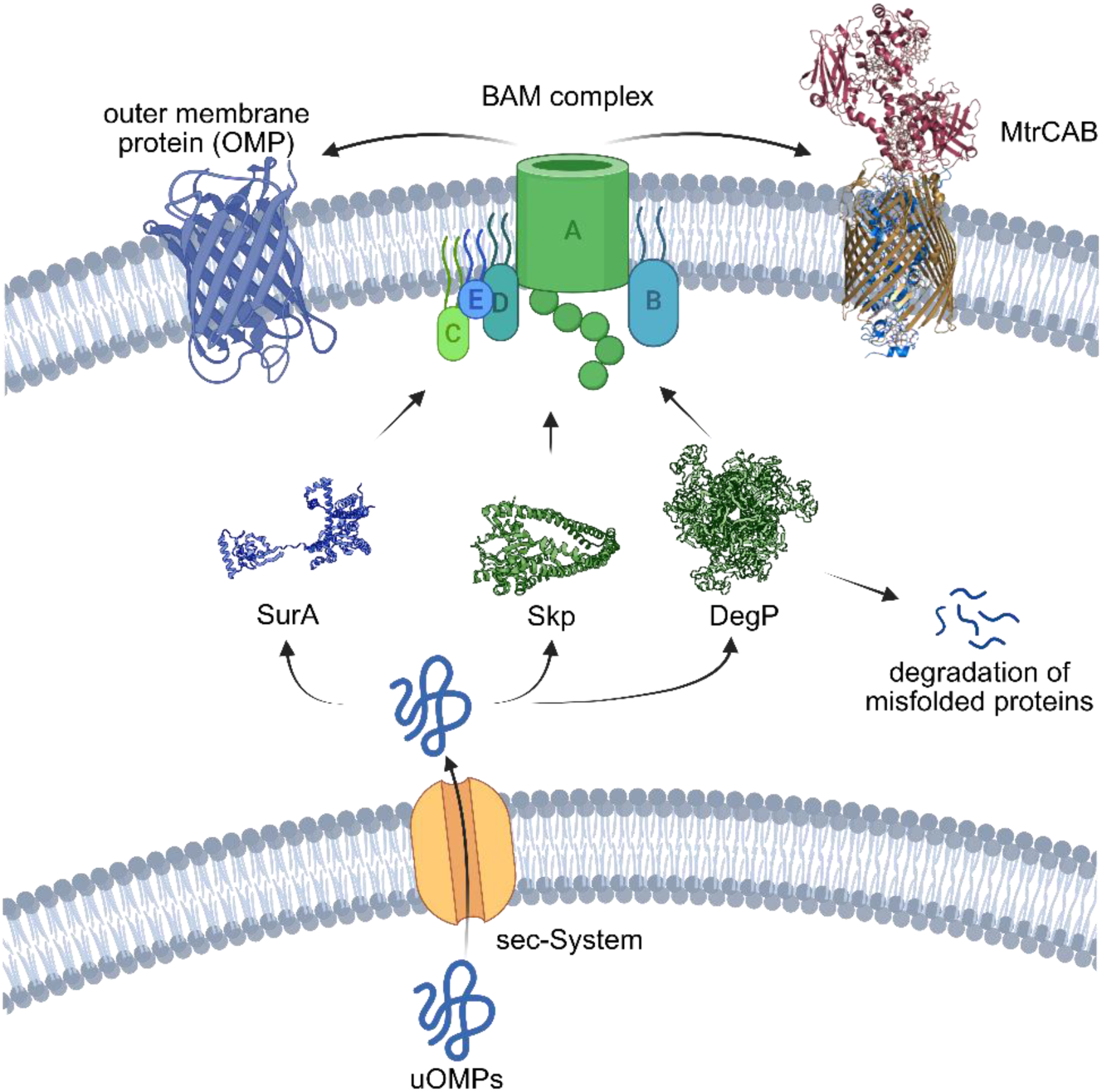
Canonical pathway of OMP transport through the periplasm. uOMPs are translocated into the periplasm via the Sec-system. The uOMP is then escorted through the periplasm by the chaperones. The two main pathways for this process, via SurA or Skp/DegP, are shown in blue and green. The uOMPs are finally released to the BAM complex which is responsible for OMP integration into the outer membrane. Crystal structures of SurA (PDB: 1M5Y ^54^), Skp (PDB: 1U2M ^55^) and DegP (PDB: 1KY9 ^56^) originate from *E. coli* K-12. The crystal structure of the MtrCAB complex (PDB: 6R2Q ^34^) is derived from *S. baltica* OS0185.

The extracellular electron transfer (EET) systems of metal-reducing bacteria provide an interesting model for investigating potentially novel aspects of OMP biogenesis. This respiratory process, which allows microorganisms to access insoluble or membrane-impermeable electron acceptors, has profound implications for global biogeochemical cycles, particularly in iron cycling^15–19^. Upon microbial reduction, ferric iron dissolves into ferrous iron, becoming available as a trace element, while simultaneously initiating abiotic geochemical reactions that reshape the composition of soils and sediments. *Shewanella oneidensis* MR-1 serves as a model organism for the study of metal reduction^20^. Metal respiration in this organism depends on the MtrCAB complex, which forms an electron conduit in the outer membrane^21,22^. This complex consists of two *c*-type cytochromes located on opposite sides of the outer membrane (periplasmic MtrA and extracellular MtrC) connected by the β-barrel protein MtrB, which facilitates their interaction^17^. Notably, gene clusters encoding MtrA-like decaheme cytochromes followed by MtrB-like β-barrel proteins are widely distributed in microbial genomes, suggesting that this arrangement represents a conserved strategy for transmembrane electron transport in both metal-reducing and metal-oxidizing bacteria^23,24^.

Despite the extensive characterization of the functional role of the MtrCAB complex, the mechanisms governing its assembly remain incompletely understood. While all three components enter the periplasm via the general secretion pathway^25,26^, with MtrA and MtrC subsequently acquiring heme cofactors via the *c*-type cytochrome maturation machinery^27–29^ and MtrC acquiring a lipid anchor^26,30^, the process by which MtrB is transported across the periplasm and correctly incorporated into the outer membrane has remained enigmatic. Interestingly, deletion of MtrA prevents proper localization of MtrB^31^, and a strep-tagged version of MtrB is degraded in the absence of MtrA, suggesting a possible non-canonical relationship between these proteins. The degradation could be overcome by a deletion of DegP^32^. However, heterologous expression studies show that even with MtrA co-expression, MtrB fails to achieve its native folded state in *E. coli*^33^, suggesting additional complexity in the assembly process. Recent structural analysis of the MtrCAB complex from *Shewanella baltica* OS185 revealed that MtrA is almost completely encased within MtrB^34^ creating a molecular wire in which MtrA is isolated from the hydrophobic membrane interior. While this structure elegantly explains the functional arrangement of the complex, it does not address the question of how MtrB traverses the periplasm prior to membrane integration.

In this study, the long-standing paradigm of obligate chaperone dependence for OMP biogenesis is challenged. Through systematic analysis using deletion mutants, conditional chaperone-null systems and quantitative proteomics, we show that MtrA functions not only as an electron transfer component but also as a highly specific chaperone for MtrB. Using Spearman correlation coefficients, we revealed that while MtrB abundance is significantly correlated with the BAM complex, it shows no significant correlation with canonical chaperones. Furthermore, we identified three additional OMPs that appear to use non-canonical trafficking mechanisms, suggesting a higher diversity in periplasmic protein trafficking strategies than previously recognized. Our findings reveal a novel specialized chaperone mechanism that has evolved in response to functional necessity, with MtrA serving the dual purpose of electron transfer and protein maturation partner for MtrB. This discovery not only advances our understanding of extracellular electron transfer systems, but fundamentally expands the paradigm of OMP biogenesis in Gram-negative bacteria, with potential implications for synthetic biology applications and our broader understanding of protein folding and transport mechanisms.

## Results

### MtrB is transported through the periplasm of E. coli by the secondary chaperone pathway

It has been demonstrated through previous and current studies that the production of a strep-tagged version of the β-barrel protein MtrB is not possible in *E. coli* when heterologously expressed, in the absence of the periplasmic decaheme cytochrome MtrA (Fig. S1). This failure in expression can be attributed to DegP-mediated degradation of strep-tagged MtrB within the periplasm^32^. The necessity of MtrA can be attributed to two potential factors. Firstly, MtrA may possess a chaperone-like function, thereby facilitating the transfer of unfolded MtrB through the periplasm. Secondly, MtrA could interact with DegP, which could in turn prevent the degradation of MtrB. Unexpectedly, this study revealed that the wild-type version of MtrB (lacking the strep-tag) can be produced in *E. coli* even in the absence of MtrA (Fig. S1). Consequently, the strep-tag sequence appears to facilitate the DegP-catalyzed degradation of MtrB in *E. coli* in the absence of MtrA. The capacity to detect wild-type MtrB using a specific antibody ^35^ facilitated the study of the interaction of MtrB with the canonical chaperones in *E. coli*. We conducted a western blot analysis with single deletion mutants of the genes encoding the chaperones SurA, Skp, and DegP. This analysis was performed to elucidate whether one of these chaperones escorts wild-type MtrB through the periplasm and releases the client to the BAM complex. However, single deletion mutants did not demonstrate any observable effects, as MtrB was detected in all outer membrane fractions (Fig. 2A). It is noteworthy that the absence of Skp resulted in the fragmentation of MtrB, which prompted further investigation into whether the Skp/DegP pathway is the primary strategy for MtrB escort through the periplasm in *E. coli*. Consequently, a Δ*skp*Δ*degP* double deletion mutant was produced and an attempt was made to express MtrB in the absence of the secondary chaperone pathway of *E. coli*. As illustrated in Figure 2A, the deletion of both genes appears to result in the degradation of MtrB. Therefore, MtrB is predominantly transported through the *E. coli* periplasm via the Skp/DegP pathway.

**Figure 2.**
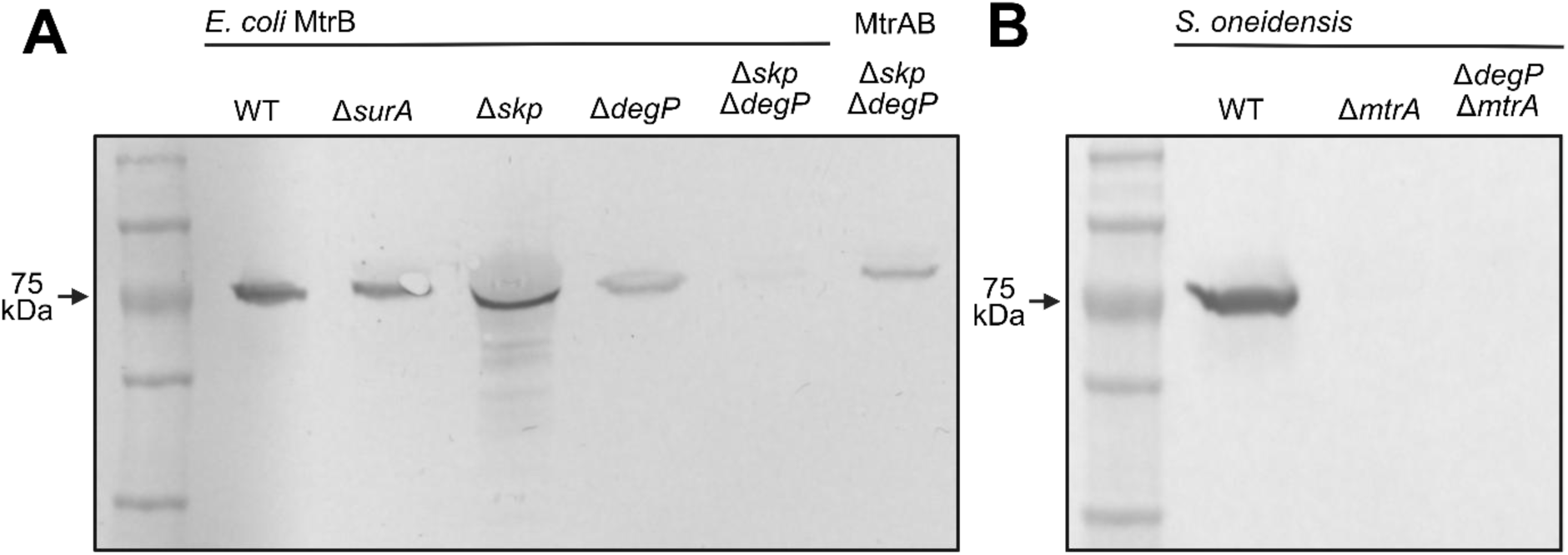
Western blot analysis of membrane fractions using an antibody specific for MtrB^35^. **A)** MtrB expression in different deletion mutants in the chaperone system of *E. coli*. pBAD-based expression of MtrB was detectable in either of the tested mutants Δ*surA*, Δ*skp* and Δ*degP*. Deletion of *skp* as well as *degP* (Δ*degP* Δ*skp*) leads to degradation of MtrB, while additional expression of MtrA restores MtrB maturation (MtrAB Δ*skp*Δ*degP*). **B)** MtrB expression in different deletion mutants of *S. oneidensis*. MtrB is only detectable when MtrA is co-expressed, additional deletion of the protease and chaperone DegP did not rescue the phenotype.

### MtrA and Skp/DegP have a redundant role for MtrB biogenesis in E. coli

It was hypothesized that MtrA acts as a specific chaperone for MtrB^32^. To investigate this further, we explored whether co-expression of MtrA in an *E. coli* Δ*skp*Δ*degP* double deletion mutant could potentially rescue the phenotype. The corresponding western blot revealed that MtrA co-expression is sufficient to compensate for the loss of Skp and DegP (Fig. 2A). In this strain, MtrB can be detected to a similar extent in the double mutant compared to the wild-type or single chaperone deletion mutants. This suggests that MtrA and Skp/DegP may have a similar function for MtrB maturation. It is interesting to note that SurA does not appear to be involved in this process, as *skp/degP* deletion results in a complete loss of the signal in western blots. However, recent findings by Philipp et al. suggest that MtrA co-expression alone may not fully replicate the same folding state observed in *S. oneidensis*^33^. Therefore, we set out to investigate whether the chaperone role of MtrA in *E. coli* resembles its native role in *S. oneidensis*.

### MtrA is crucial to prevent degradation of MtrB in S. oneidensis

Schicklberger et al. demonstrated that in *S. oneidensis*, a strep-tagged version of MtrB is degraded in the absence of MtrA. Conversely, it is unaffected when *degP* (SO_3942) is additionally deleted^32^. This finding is consistent with the observation that MtrB_strep_ can only be detected in conjunction with MtrA during co-expression in *E. coli* (Fig. S1). However, our previous findings indicated that this effect can be directly attributed to the strep-tag, rendering it more susceptible to degradation (Fig. S1). Therefore, we investigated the potential for the detection of native MtrB in membrane fractions of a *S. oneidensis* Δ*mtrA* mutant. In contrast to *E. coli*, where MtrB is readily identifiable, its presence was not detected in the absence of MtrA in *S. oneidensis*. This observation indicates that MtrB’s reliance on MtrA is more pronounced in *S. oneidensis* compared to *E. coli*. This phenomenon remained constant even in the context of testing the Δ*mtrA*Δ*degP* double deletion mutant (Fig. 2B). Consequently, the absence of DegP does not counterbalance the absence of MtrA^32^. This finding indicates that DegP plays a role in MtrB degradation exclusively in the presence of a strep-tag attached to the uOMP in *S. oneidensis*. The findings indicate that SurA appears to be dispensable for MtrB chaperoning, whereas Skp/DegP can escort it to the outer membrane in *E. coli*, though not in *S. oneidensis*. At this juncture, the potential involvement of Skp or SurA to aid in MtrB transport through the periplasm in *S. oneidensis* cannot be excluded, despite the demonstration of a more pronounced dependency on MtrA (Fig. 2B). Hence, a chaperone-null mutant was necessary to fully understand the role of the canonical chaperones in this process.

### Construction of a S. oneidensis conditional chaperone-null mutant

While the construction of a *degP* single deletion mutant was technically feasible, the introduction of a second chaperone deletion appeared to be lethal in *S. oneidensis*. Therefore, the initial step was to introduce inducible versions of *skp* and *surA*, thereby enabling deletion of the native loci in subsequent steps. Consequently, the tac promoter^36^ was selected to ensure sufficient and inducible expression. In order to circumvent potential homologous recombinations between the native loci and the introduced synthetic *skp-surA*, the new genes underwent modifications to reduce sequence similarities while maintaining the overall codon usage. To this end, the frequency of each codon for every amino acid in the respective wild type *surA* and *skp* sequences was calculated. Subsequently, a random exchange of codons was performed, replacing them with codons that are similarly abundant. An overview of the used codons is provided in table S1. The modified versions of *skp* and *surA* exhibit no more than 16 consecutive identical nucleotides in common with the native versions, and their overall sequence identity ranges from 69.9% to 74.3%, as for the *surA* and *skp* gene, respectively.

The *surA-skp* operon was integrated into the *S. oneidensis* genome in conjunction with the *lacI* gene. The introduction of the modified chaperones was concomitant with the deletion of *degP*, and all subsequent steps were carried out in the presence of 1 mM IPTG to facilitate expression of the modified chaperones. Subsequently, the *surA* gene was successfully deleted through the implementation of the standard homologous recombination protocol. Nevertheless, the process of *skp* deletion was found to be impossible. Despite multiple attempts, only revertants were obtained. It was hypothesized that this phenomenon is attributable to polar effects exerted by the essential gene *bamA*, which is located in close proximity to *skp*. Consequently, a deactivated CAS enzyme was utilized in conjunction with a deaminase^37^ to insert a stop codon within the *skp* sequence (p.Gln115*, encompassing a total of 168 amino acids), thereby resulting in a truncated Skp in the *S. oneidensis* Δ*chaperone* (Δ*degP*::P_tac_-*skp_cm__surA_cm_* Δ*surA skp\**) strain (Fig. 3A). In the absence of induction of *surA* and *skp*, this strain is devoid of the chaperones known to be involved in OMP biogenesis. Consequently, it offers a suitable model for direct study of the role of the canonical chaperones and the interaction between MtrA and MtrB in the periplasm (Fig. 3B).

**Figure 3.**
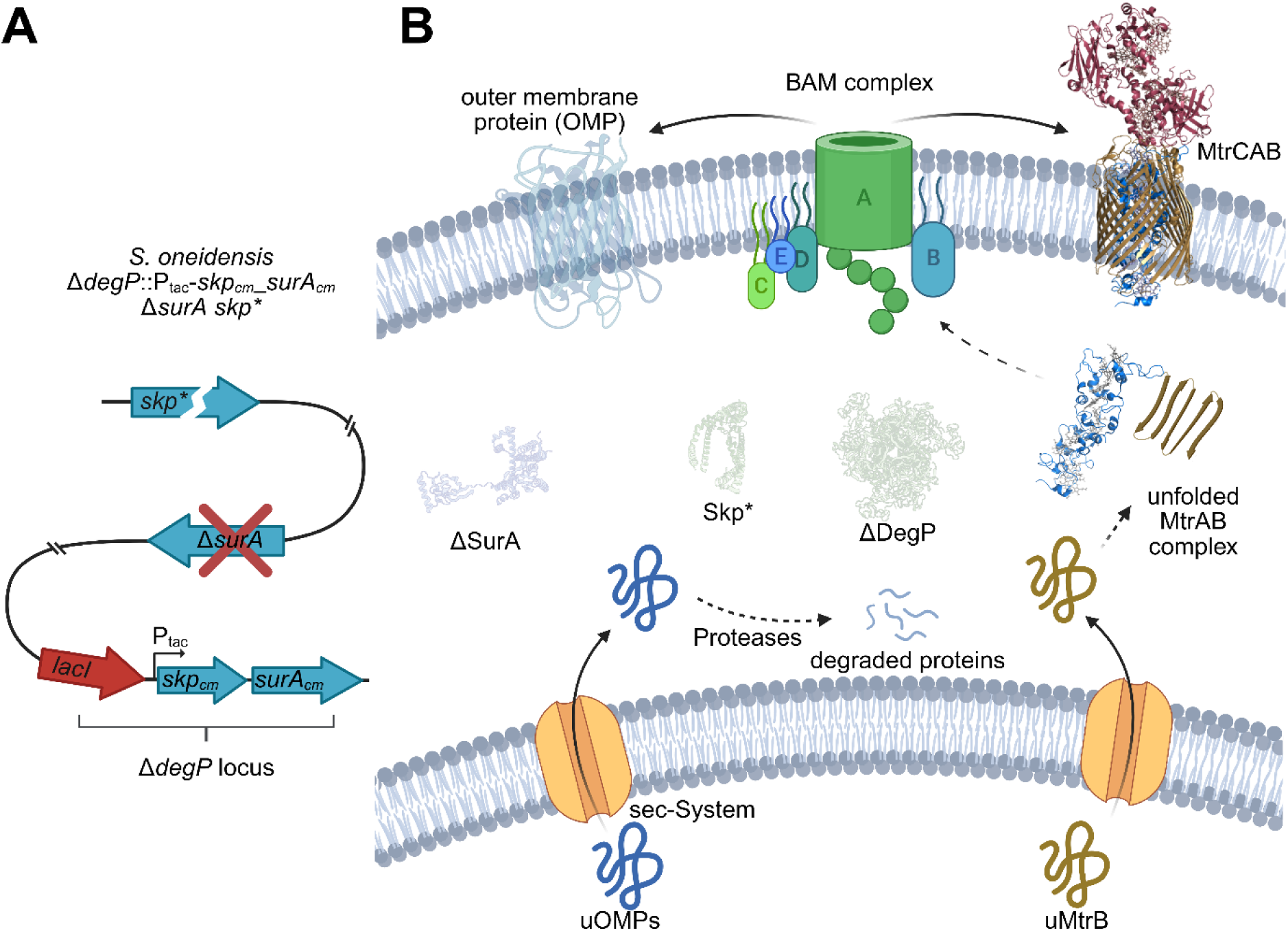
Genotype and expected phenotype of the conditional *S. oneidensis* Δ*chaperone* mutant. **A)** Codon modified (cm) versions of *skp* and *surA* are introduced into the degP locus and expressed upon induction of the tac promoter. SurA was deleted via homologous recombination while a stop codon was introduced to truncate Skp. **B)** Expected phenotype of the condtional mutant without induction. Depletion of the chaperones SurA, Skp and DegP and consequently the unfolded outer membrane proteins (uOMPs) is represented through higher transparency. MtrB could take an MtrA-assisted route through the periplasm and is incorporated into the outer membrane via the BAM complex. Crystal structures of SurA (PDB: 1M5Y^54^), Skp (PDB: 1U2M^55^) and DegP (PDB: 1KY9^56^) originate from *E. coli* K-12. The crystal structure of the MtrCAB complex (PDB: 6R2Q ^34^) is derived from *S. baltica* OS0185.

### MtrA is the only chaperone for MtrB in a conditional chaperone null mutant

To assess the impact of chaperone depletion, we cultivated the induced *S. oneidensis Δchaperone* strain in oxic LB media for an overnight period. Following the transfer to new medium, one replicate was supplemented with IPTG to promote chaperone expression, while another replicate was cultivated in the absence of IPTG, resulting in chaperone expression being absent in this replicate (see Figure 4A). A sample was collected for protein analysis as soon as the no-chaperone cultures exhibited growth deficiencies (after 9 hours or 4 generations, Fig. 4A t1), suggesting that the chaperones were diluted to such an extent that uOMPs were no longer handed over to the BAM complex. A second sample was collected subsequent to reaching the stationary phase (after 16 hours, Fig. 4A t2). Western blot analysis of samples obtained at t1 and t2 revealed that MtrB remains present in the membrane fraction at comparable levels in the presence and absence of the canonical chaperones (Fig. 4B). Western blots carried out with an anti-Omp35 antibody functioned as a positive control for chaperone depletion. It is noteworthy that Omp35 is listed among the top 5% of the most abundant proteins in *S. oneidensis* according to the database paxDB^38^, thus serving as the positive control for a canonical ß-barrel protein. In fact, the strains lacking chaperone induction exhibited a considerably weaker band for Omp35, indicating that this ß-barrel protein undergoes rapid degradation in the absence of the canonical chaperones. This finding corroborates the efficacy of this approach in elucidating the role of SurA and Skp/DegP in *S. oneidensis* (Fig. 4C). The severe growth defect exhibited by the conditional chaperone-null mutant suggests that *S. oneidensis* is incapable of compensating for the simultaneous loss of SurA and Skp/DegP. Nevertheless, MtrB production is not affected. However, the assessment conducted thus far has been limited to a qualitative evaluation of the production of two ß-barrel proteins. In order to facilitate a more quantitative evaluation of the collective impact of chaperones on uOMP transport, a more comprehensive approach was implemented in a subsequent phase of the study.

**Figure 4.**
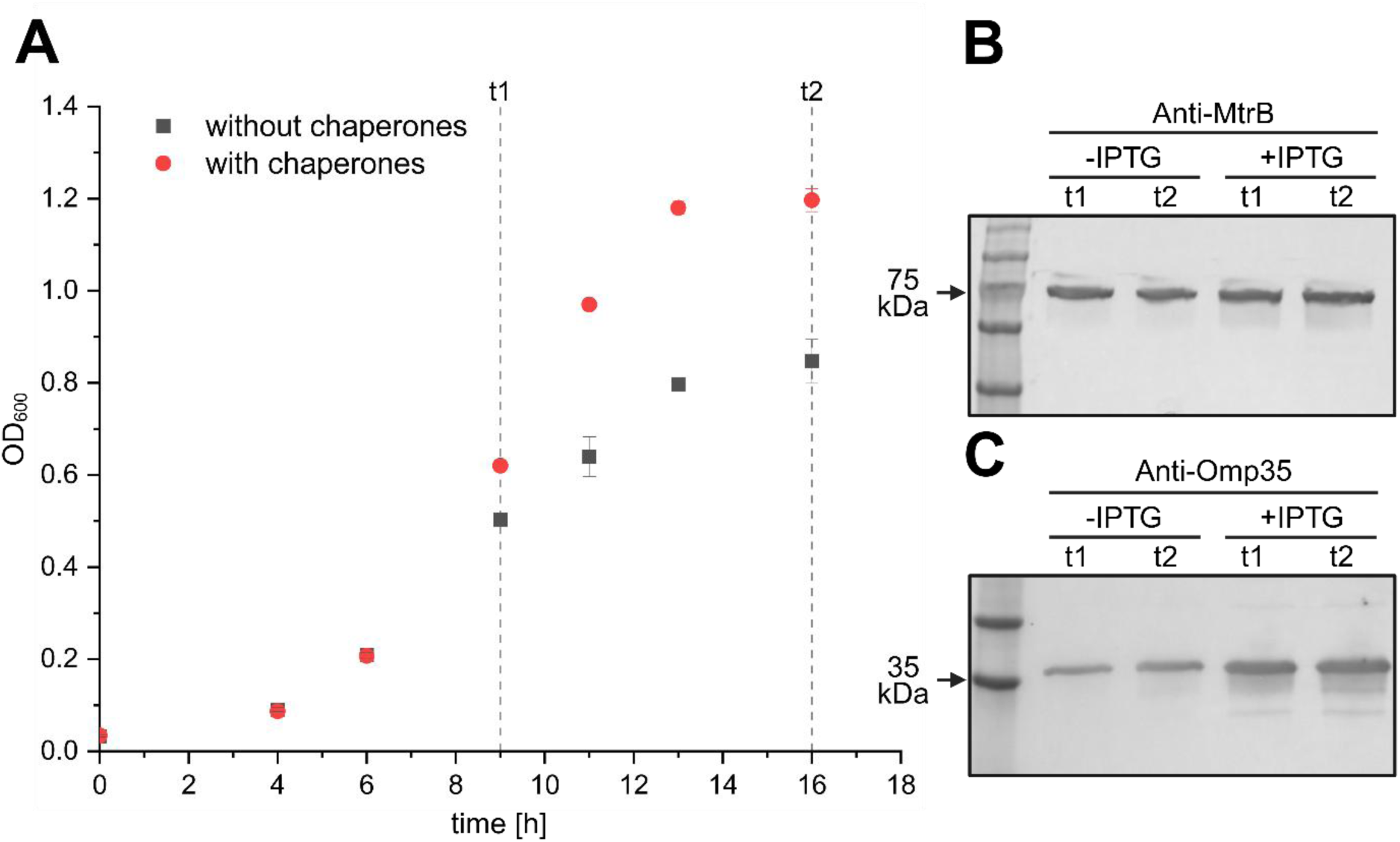
Growth and Immunoblotting of the conditional *S. oneidensis* Δ*chaperone* mutant. **A)** Growth curve with (red circles) and without (black squares) induction of *surA* and *skp*. The first sample was taken after four cell divisions (t1), when growth began to differ between the conditions. The second sample was taken after reaching the stationary phase (t2). Data points represent the average of three independent replicates, error bars indicate the standard deviation. **B)** Western blot analysis of 30 µg membrane fractions using an anti-MtrB antibody ^35^ without (-IPTG) and with induction (+IPTG). MtrB is detactable regardless of the presence of the canonical chaperones SurA, Skp and DegP in *S. oneidensis*. **C)** Western blot analysis of 2.5 µg membrane fractions using an anti-Omp35 antibody without (-IPTG) and with induction (+IPTG). The signal is weaker when the chaperones are not induced.

### Deciphering of SurA and Skp/DegQ dependency on uOMPs in S. oneidensis

A quantitative proteome analysis of the soluble and membrane fractions was conducted, and the difference between the induced and non-induced samples at t1 and t2 was analyzed. As anticipated, DegP was not detected in the proteome, given that the corresponding gene had been deleted and was not part of the synthetic, inducible chaperone cluster. The quantification of SurA was found to be consistent across all samples, and it was identified as one of the most differentially regulated proteins within the proteome. Specifically, SurA exhibited the highest negative log_2_ fold change in membrane fractions at timepoints t1 and t2, the most significant negative log_2_ fold change in soluble fractions at t2, and the second most significant negative log_2_ fold change at t1. These findings once again demonstrate the applicability of the conditional chaperone null mutant. The presence of Skp in the proteome was identified; however, its quantification was only possible in the induced samples and not in the uninduced samples. This finding serves to demonstrate that the chaperones are rapidly diluted by growth without expression.

In order to comprehend the impact of SurA and Skp/DegP on the stability of uOMPs, it was imperative to first identify all potential ß-barrel proteins within the *S. oneidensis* Δ*chaperone* proteome. A DeepTMHMM^39^ search was conducted on all proteins detected in the membrane fraction to identify produced ß-barrel proteins. A total of 52 potential ß-barrel proteins were identified (see Table 1). This list was then manually curated through comparison with AlphaFold structures and exclusion of proteins with no more than four ß-sheets (AggA; TolC; an ABC-type efflux system secretin component SO_4090; a CzcC family protein SO_0518). These proteins have been observed to form trimeric ß-barrels that are not dependent on periplasmic chaperones^40–42^. A total of 48 ß-barrel proteins remained to be analyzed; however, 12 of these were not abundant enough to allow for quantification. Consequently, 36 ß-barrel proteins could be used for statistical analysis. A comparison was made between the non-induced cells and the induced cells. The log_2_ fold changes were subsequently calculated, with the respective p-values. At time point t1, 26 of the 36 proteins demonstrated significant downregulation (log_2_ fold change < −1, p-value < 0.05) in the samples lacking chaperones. Notably, one ß-barrel protein (SO_4422) exhibited a decrease in expression, although this variation did not attain statistical significance (log_2_ fold change < −1, p-value > 0.05). Surprisingly, one of the genes (SO_3810) demonstrated a substantial increase in expression (log_2_ fold change = 1.4, p-value < 0.05), while eight genes, including MtrB, exhibited no significant regulation (log_2_ fold change > −1 and < 1). This finding indicates that these genes are not influenced by the chaperone depletion (Fig. 5A). A subsequent analysis revealed that these eight proteins could be divided into two distinct clusters. The initial group comprises three OMPs (YfaZ, SO_0934, SO_3931), exhibiting a minimal and non-statistically significant log_2_ fold change ranging from −0.2 to 0.2. The second group (MtrB, SO_3905, SO_3545, SO_3099, SO_2736) exhibits a set of proteins that undergo only a negligible change in abundance, with log_2_ fold changes between −1 and 1, when comparing not induced cells vs. induced cells. However, for two of these proteins (SO_2736; SO_3905), the observed deviation in concentration exceeded the established threshold for statistical significance, precluding the ability to ascertain any meaningful trends or variations. In the stationary phase, the minor discrepancy between the uninduced and induced samples for the second group was further mitigated for MtrB, SO_3545, and SO_2736. Conversely, the log_2_ fold change remained constant for SO_3905 and SO_3099 (see Table S2). Consequently, the outcomes of western blotting are substantiated by quantitative proteomic analysis. As illustrated in Figures 5A and 5B and Table S2, the concentration of MtrB remains nearly constant. Conversely, a significant array of ß-barrel proteins undergoes a reduction in concentration, resulting from the downregulation of periplasmic chaperones.

**Figure 5.**
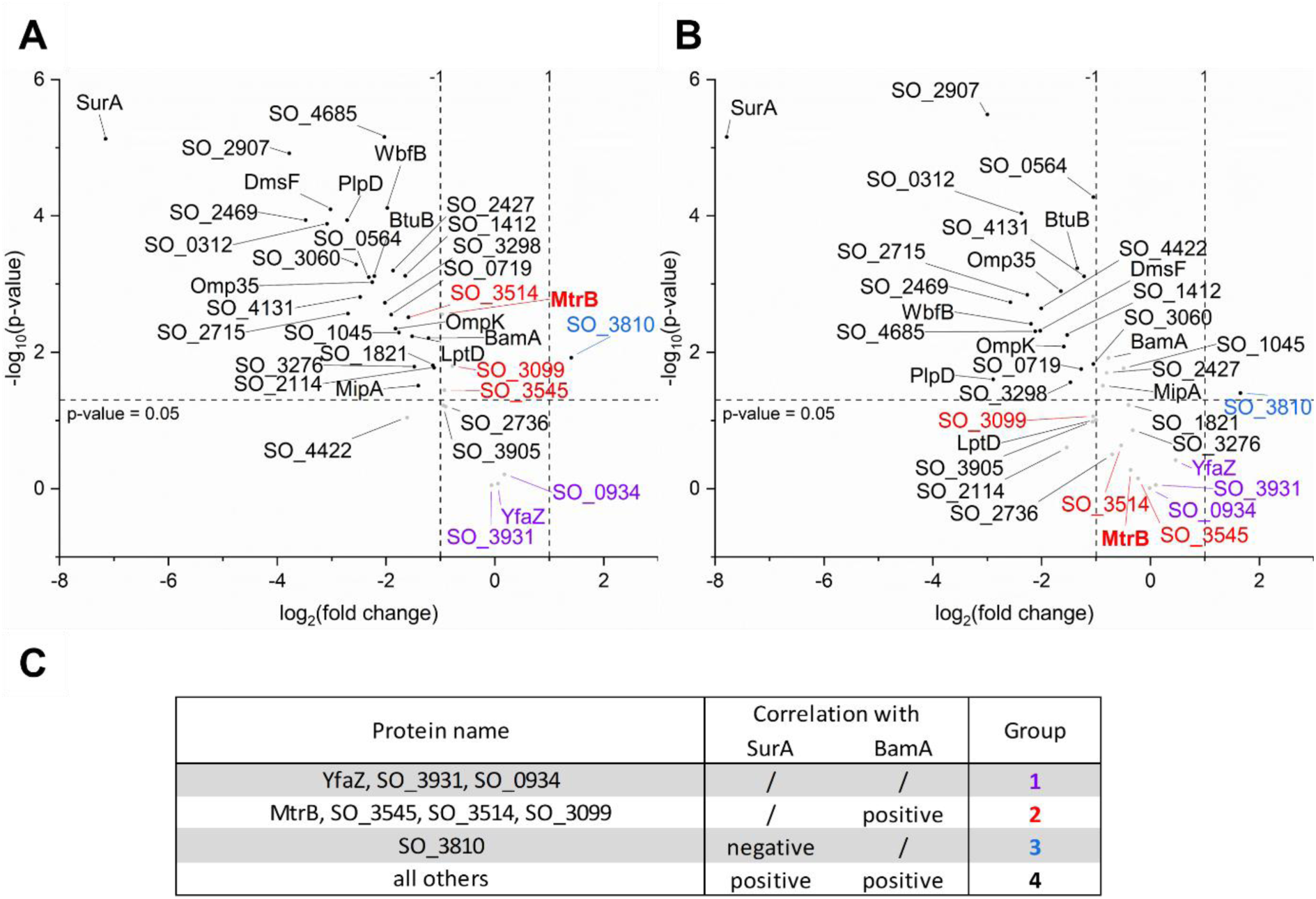
Volcano plots of ß-barrel protein abundance in membrane fractions after growth without induction of the chaperones vs. with induction. Black dots represent a significant log_2_(fold change) (log_2_ fc < −1 or > 1, p-value < 0.05), grey dots represent no significant change in **A)** the exponential phase or **B)** stationary phase. The vertical dashed lines represent the threshold for log_2_(fold change) while the horizontal dashed line represents the p-value threshold. Log_2_(fold change) and p-values are calculated from independent triplicates of each condition. **C)** Spearman correlation of the ß-barrel proteins with SurA or BamA, respectively. Coefficients are calculated from all replicates of the two time points and two conditions. A slash represents no significant correlation (p-value > 0.05) while positive and negative indicates significant positive or negative correlations (p-value < 0.05). The font color of the four groups is also reflected in the volcano plots.

**Table 1.**
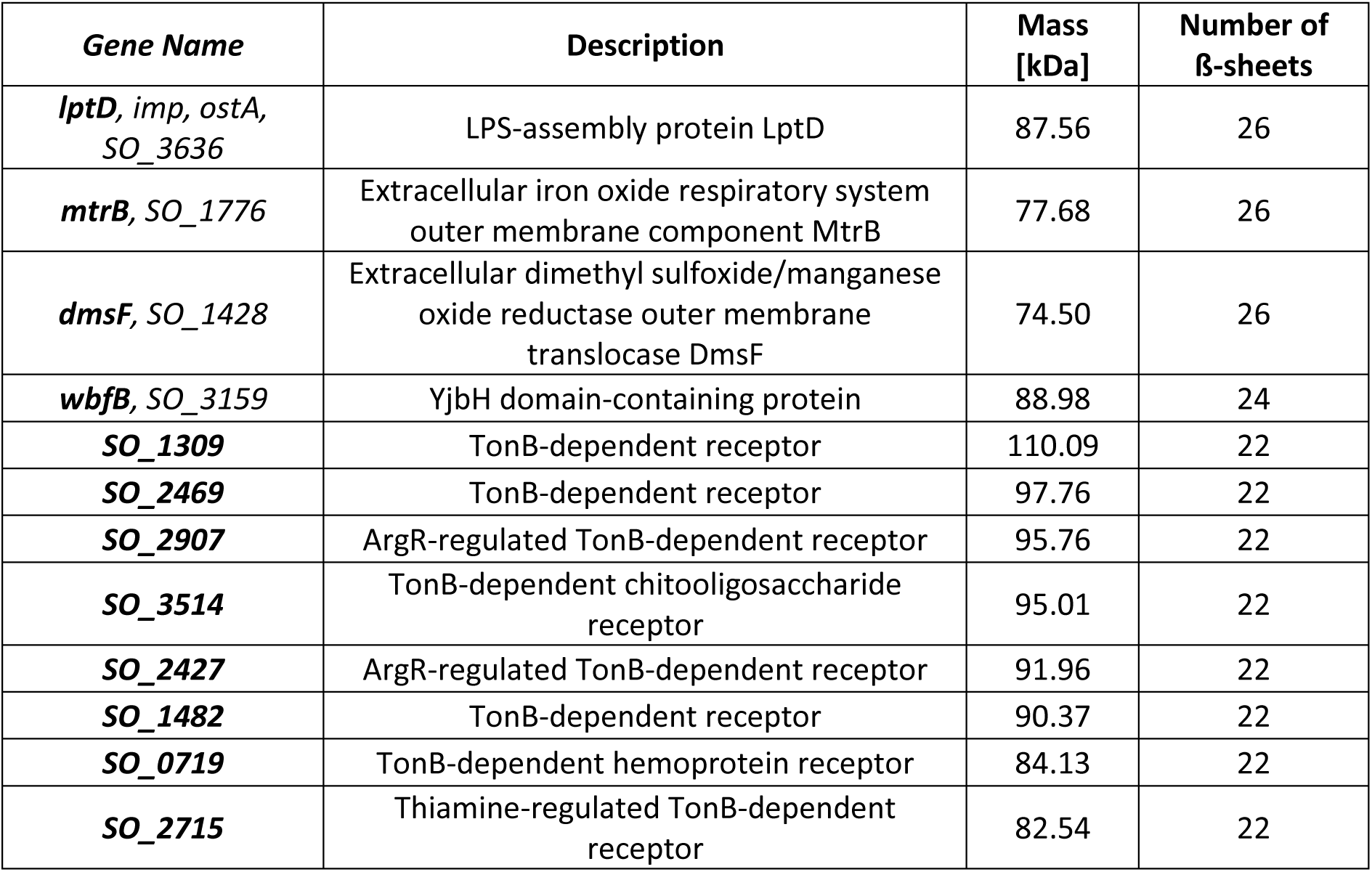
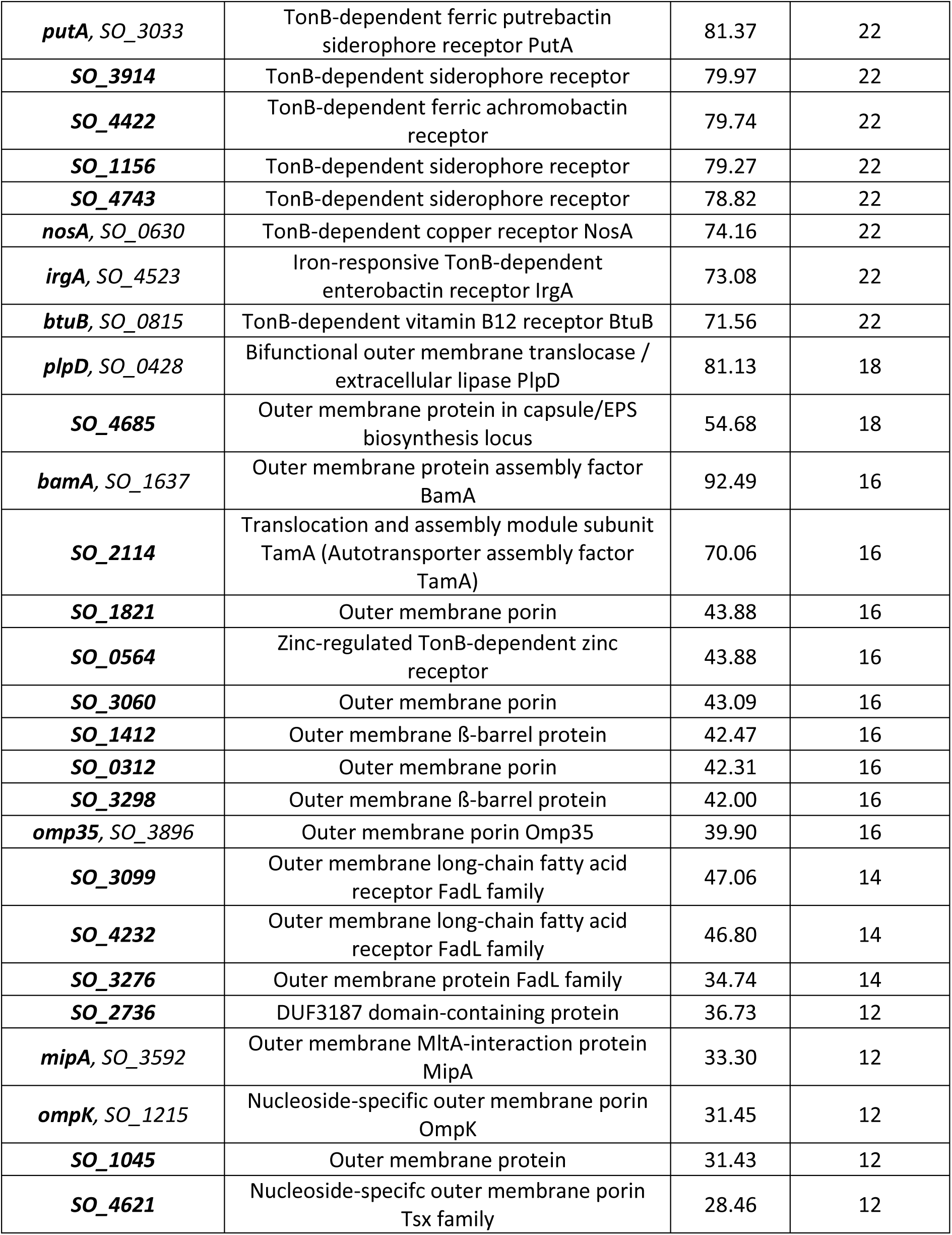
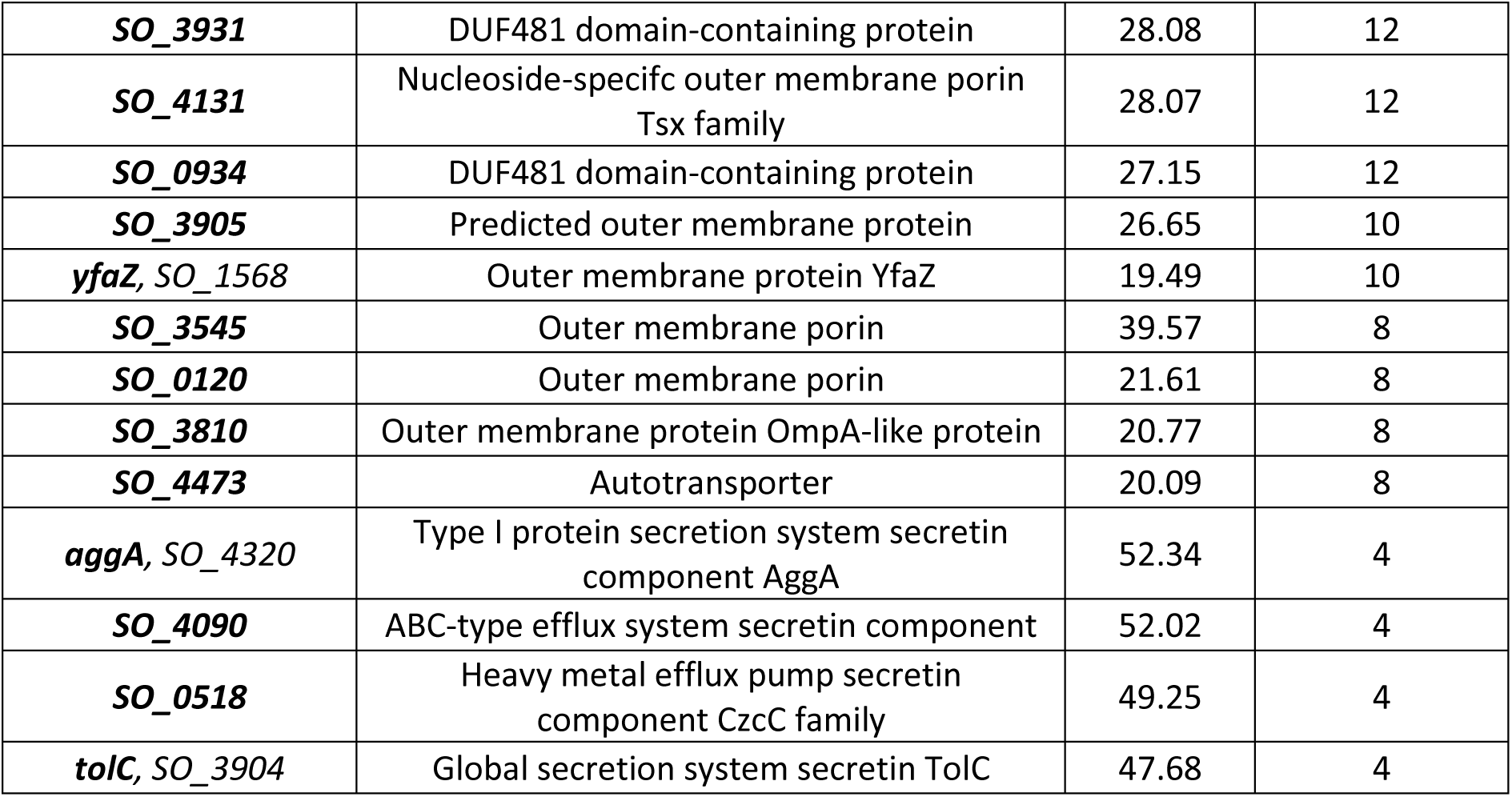
Overview of the predicted ß-barrel proteins in the membrane fractions of the proteome with the respective size in kDa and number of ß-sheets.

### The small concentration change of MtrB under chaperone downregulation conditions correlates with BAM complex but not with SurA abundance

Apart from the chaperones SurA and Skp/DegP, the BAM complex plays an essential role in the process of protein assembly into the outer membrane. BamA, an integral member of the BAM complex, is a β-barrel protein. Its abundance is significantly reduced in cells with reduced chaperone expression in the exponential phase (t1, log_2_ fold change = −1.22, p-value = 0.0062). However, its abundance is not significantly different in the stationary phase (t2, log_2_ fold change = −0.77, p = 0.0121). Therefore, it was sought to determine whether the primary effect of chaperone depletion was responsible for uOMP degradation or if the secondary effect of a reduced BAM complex availability was the main reason. To this end, Spearman correlation coefficients were calculated to test the hypothesis of a significant (p-value < 0.05) correlation with SurA and/or BamA. A significant positive correlation was observed between 28 OMPs and both SurA and BamA, suggesting that the maturation pathway for these 28 proteins follows the canonical pattern (Fig. 5C, Group 4). SO_3810 demonstrated a significant negative correlation with SurA but not with BamA (Fig. 5C, Group 3). The three OMPs that demonstrated complete resistance to the effects of chaperone depletion (SO_3931, SO_0934, and YfaZ) exhibited no correlation with SurA or BamA (Fig. 5C, Group 1).

A correlation was identified between four OMPs (SO_3099, SO_3514, SO_3545, and MtrB) and BamA, but not SurA (Fig. 5C, Group 2). These findings suggest that the assembly of these proteins into the outer membrane occurs via the canonical BAM complex, while they do not rely on the canonical chaperons in the periplasm. The calculation of correlation coefficients between Skp and ß-barrel proteins was not feasible due to the low abundance of Skp in the samples lacking chaperones. However, the data suggest that the observed variation in MtrB quantity between samples with and without chaperones in exponential phase (t1) is a consequence of reduced BamA quantity. Consequently, the observed difference can only be attributed indirectly to the loss of chaperones. This finding is consistent with the results of the western blots, which demonstrated complete degradation of MtrB in the absence of MtrA (Fig. 2B) in *S. oneidensis*, while the loss of chaperones had no effect (Fig. 4B).

## Discussion

The present study employed a combination of *E. coli*- and *S. oneidensis*-based experiments to elucidate the function of canonical chaperones, as well as MtrA, in the context of MtrB biogenesis. We used chaperone deletion mutants in *E. coli* as well as in *S. oneidensis* to investigate the following: first, whether MtrA can indeed guide MtrB through the periplasm; and second, whether this is only a rescue process if MtrB falls off the main chaperone guided pathway or indeed the primary pathway. Given that MtrB can be produced in *E. coli* without MtrA, it is evident that in this organism, a second pathway is capable of fulfilling the chaperone role for MtrB. Indeed, the deletion of the Skp/DegP pathways resulted in the loss of MtrB in an *E. coli* strain lacking MtrA, thereby demonstrating that Skp and DegP can both act as chaperones for MtrB (Fig. 2A). The phenotype of the double mutant can be rescued through co-expression of MtrA, thereby demonstrating, for the first time, that MtrA can substitute for the role of chaperones (Fig. 2A). To further investigate the relationship between MtrA and its client MtrB, we tested the dependency in *S. oneidensis*. In this system, MtrA is indispensable for MtrB biogenesis, and the deletion of the gene for protease DegP is unable to prevent MtrB degradation in Δ*mtrA* mutants (Fig. 2B). The results of this study demonstrate both similarities and differences to the LptD/E system in *E. coli*. It has been demonstrated that the proteins LptD and LptE can only be overexpressed in conjunction, a phenomenon that bears a striking resemblance to the requirement for MtrA in the production of MtrB in *S. oneidensis*^43^. However, the impact of the lipoprotein LptE on LptD production appears to be localized to the outer membrane following an initial interaction of LptD with the BAM complex^44^. Furthermore, LptD is guided through the periplasm by the canonical SurA-pathway^45–47^. In contrast, the results of the *E. coli* experiments suggest that the interaction of MtrA with MtrB already commences during the transfer of MtrB through the periplasm. This is due to the fact that MtrA itself can replace Skp/DegP and fulfill a chaperone function for transport through the periplasm (Fig. 2A).

We generated a conditional *S. oneidensis chaperone* mutant devoid of all canonical chaperones to ascertain whether MtrA is the sole chaperone or merely assists SurA or Skp/DegP in the native organism. This mutant produced a similar band for MtrB in western blots of membrane fractions, while the control protein, Omp35, was degraded (Fig. 4B). Furthermore, the quantitative proteome corroborated the findings of the western blot analysis, as MtrB exhibited neither substantial regulation in the exponential phase (Fig. 5A) nor in the stationary phase (Fig. 5B). As illustrated in Figure 4A, the observed growth defect, in conjunction with the finding that a significant proportion of quantifiable β-barrel proteins are substantially less abundant in samples not induced, indicates that the chaperones were diluted to a degree that they were unable to escort the majority of uOMPs to the BAM complex. However, in the stationary phase, a significantly higher number of proteins were not subject to regulation (Fig. 5B). This finding suggests that either a SOS response in the cells led to an enhanced OMP assembly compared to the exponential growth phase (Fig. 5A), or that the outer membrane disintegrates in stationary-phase cells, even in the presence of chaperones.

Experiments conducted by Philipp et al.^33^ suggest that although MtrB transport to the outer membrane is possible in *E. coli* the folding of the protein in the outer membrane might differ. This latter assumption is based on the observation that certain epitopes of the folded protein are available for antibody binding in *E. coli* but not in *S. oneidensis*, as revealed by tests with antibodies. Therefore, despite the fact that MtrB is localized in *E. coli* through its interaction with MtrA or DegP/Skp, it is possible that this version exhibits a distinct functionality compared to that of *S. oneidensis*. The precise mechanism by which the MtrAB complex is correctly incorporated remains to be elucidated. However, the quantitative proteomics approach has facilitated a more profound comprehension of the underlying interaction by calculating Spearman correlation coefficients.

The aforementioned correlations permitted the elucidation of the question of whether differential regulation of OMPs is a direct result of chaperone dilution or of a non-functional BAM complex. As illustrated in Figure 5C, Group 1 proteins appear to be unregulated, exhibiting no significant correlation with SurA or BamA. This observation suggests the possibility of a non-canonical pathway being followed by these proteins. SO_3810 exhibits a substantial negative correlation with SurA, yet no correlation with BamA is observed. The question of whether the upregulation is a direct response to SurA downregulation or merely a secondary effect remains to be elucidated. This ß-barrel protein is predicted to have an OmpA-like transmembrane domain, and the corresponding gene is apparently part of a monocistronic operon. The normalized abundance of SO_3810 remains constant in the induced cells in comparison with the oxic pre-culture, while it increases in the uninduced cells (see Table S2). Therefore, it can be concluded that the expression is a response to the chaperone depletion.

All other β-barrel proteins that were analyzed (Groups 2 and 4, Fig. 5C) exhibited a significant positive correlation with the BAM complex. This finding indicates that these proteins adhere to the standard pathway for OMP incorporation into the outer membrane. The majority of these proteins (28 of the remaining 32 ß-barrel proteins) demonstrate a significant positive correlation with SurA as well (Fig. 5C, Group 4), indicating that they traverse the periplasm via the canonical pathway. A total of four proteins, including MtrB, were found to be non-significantly correlated with SurA but were observed to be positively correlated with BamA (Fig. 5C, Group 2). A close examination reveals no obvious sequence similarities between the four proteins. Furthermore, while they are indeed ß-barrel proteins, they appear to be unrelated in terms of their functions and structures. Therefore, there is a high probability that their mechanism or periplasmic transfer underwent independent evolution.

The results with *S. oneidensis* suggest that the ability of *E. coli* Skp/DegP to guide MtrB through the periplasm might be an artefact that is not relevant under native conditions. Nevertheless, even the co-expression of MtrAB might result in the malfunctioning of the conduit in *E. coli*. The latter suggests that the incorporation of one or more additional functionalities from *S. oneidensis* may be essential for establishing effective interaction between MtrA and MtrB in *E. coli*. Nonetheless, it is conceivable that the overall structure of the outer membrane in *E. coli* differs to such a degree from that of *S. oneidensis* that proper folding becomes unfeasible, despite the presence of all requisite maturation components.

This study unveils a novel mechanism for the specific guidance of outer membrane ß-barrel proteins through the periplasmic space. The proteomic results indicate that MtrB may not be the sole protein with an individual solution, as other proteins were also unaffected by SurA and Skp depletion. The existence of additional non-canonical pathways for other ß-barrel proteins that are not essential for growth under the conditions that have been tested thus far merits further investigation.

## Material and methods

### Codon Modification of Chaperones

The codon frequency of the genes *skp* and *surA* was analyzed using the CLC Main Workbench. In this study, each codon was systematically substituted with one of the top three most prevalent codons of each amino acid. The Excel function “INDEX” was utilized to provide the array containing the three most frequently used codons of the specific amino acid. The row number was defined using the “RANDBETWEEN(1;3)” function. The corresponding Excel template is provided as supplementary information. The nucleotide sequences were checked for large identical areas and overall sequence identity using a discontiguous megablast^48^.

### Construction of knock-out mutants and expression plasmids

*E. coli* single knock-out mutants were purchased from the *E. coli* Genetic Resource Center (ECGRC, CT, USA). As previously described^49^, the *skp* gene was also deleted in the Δ*degP* strain using primers 22 and 23.

Markerless *S. oneidensis* deletion mutants were constructed using pMQ150 suicide vector as described previously^50,51^. The *degQ* gene was deleted using primers 1 – 4 in a *S. oneidensis* Δ*mtrA* strain. The DNA sequence of the repressor *lacI,* tac promoter and inducible, codon modified versions of *skp* and *surA* was ordered as cloned gene from Thermo Fisher Scientific (USA) and is given in Figure S2. For induction, 1 mM IPTG was added in all cultivation steps after introduction of the codon modified versions of *skp* and *surA* into the *degP* locus. Subsequently, *surA* was deleted using primers 14 – 17. The introduction of a premature stop codon in *skp* (p.Gln115*) was carried out as described elsewhere^37^. Briefly, a deactivated CAS enzyme was used to guide a deaminase to the desired position catalyzing the conversion of cytosine to thymine. The plasmid pCYR104 was linearized using BsaI-HF.v2 (New England Biolabs, USA). Primers 20 and 21, coding for the sgRNA to introduce a stop codon in *skp*, were resuspended in Duplex buffer (100 mM potassium acetate, 30 mM HEPES, pH 7.5), heated to 95 °C and afterwards slowly cooled to room temperature (−2 °C per minute) to obtain double stranded DNA for isothermal *in vitro* assembly. Plasmid curing in the mutant was carried out through overnight incubation without antibiotics followed by screening for kanamycin sensitive colonies.

The expression plasmid pBAD202 was linearized using the restriction enzymes NcoI-HF and PmeI (New England Biolabs, USA). All primers used are listed in table 2. All linearized vectors for genomic modifications and overexpression studies were assembled with inserts through isothermal *in vitro* assembly as described by Gibson et al.^52^.

**Table 2.**
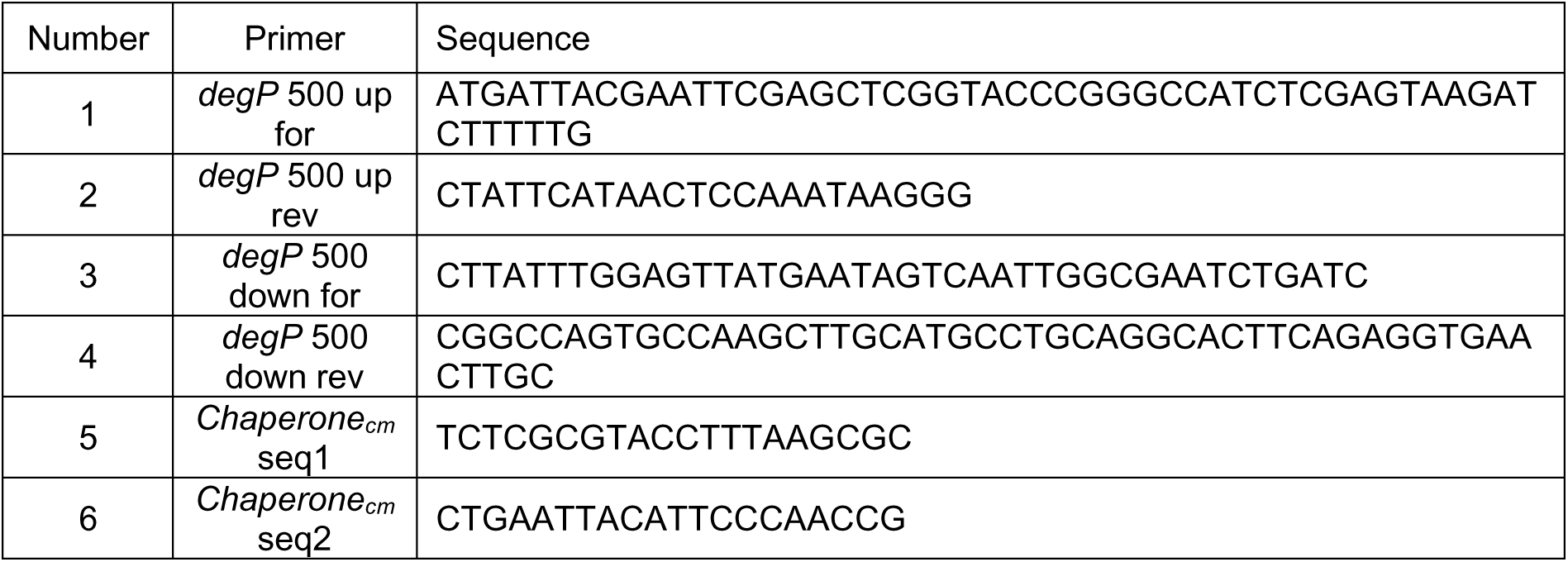
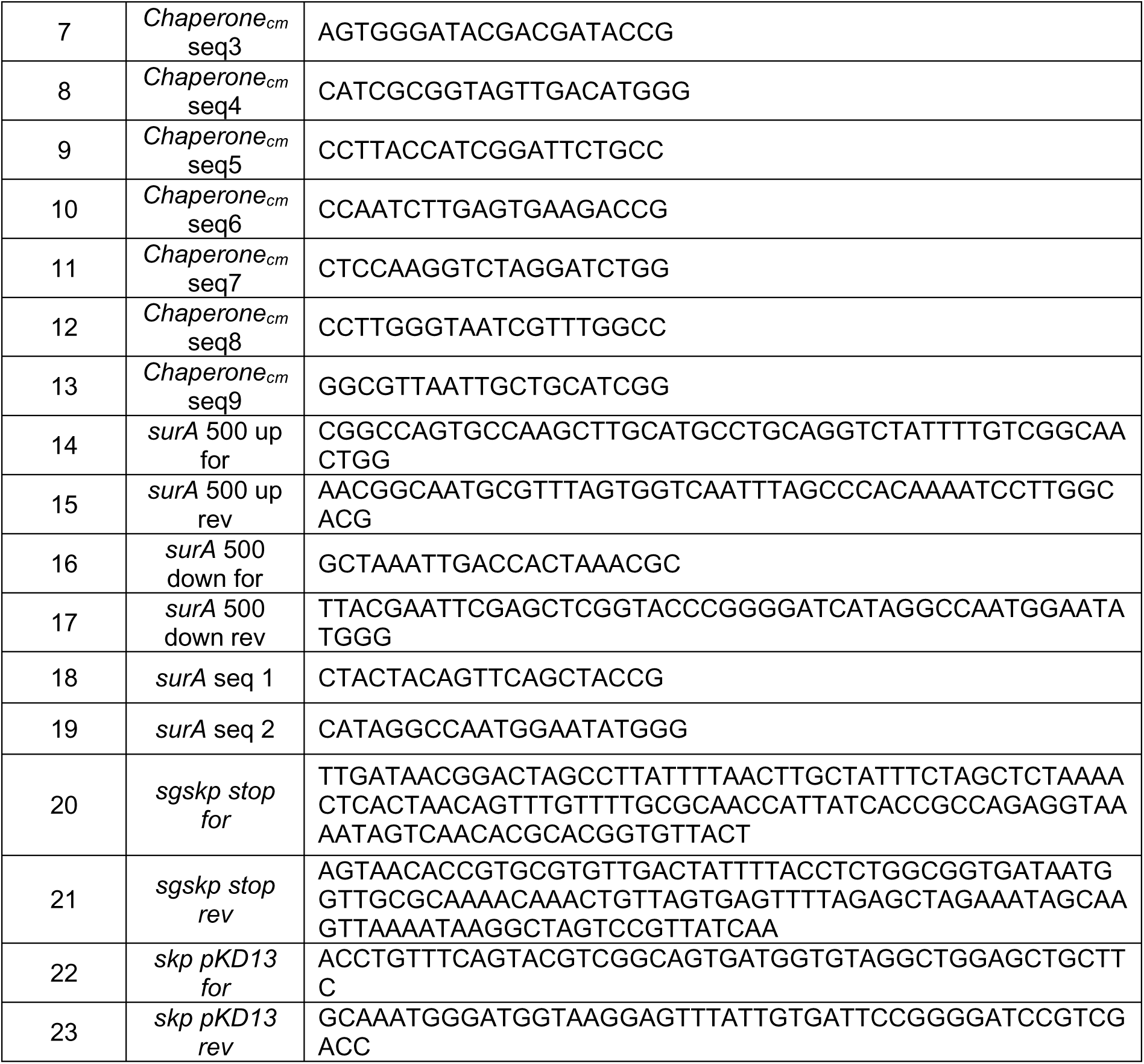
Primer*s*, used in this study.

All insertions and deletions were confirmed via Sanger sequencing at Eurofins Genomics (Luxembourg) and subsequent analysis using Benchling (CA, USA).

### Bacterial strains and culture condition

The bacterial strains used in this study are listed in table 3. The cultures were grown aerobically in lysogeny broth medium (LB). Anaerobic *S. oneidensis* cultures were grown in LB media at pH 7.4 buffered with 50 mM HEPES. Lactate (50 mM) and fumarate (100 mM) were added as additional electron donor and electron acceptor, respectively. Anaerobic *E. coli* cultures were grown in LB media at pH 7.4 buffered with 50 mM HEPES. Glycerol (50 mM) and fumarate (100 mM) were added as additional electron donor and electron acceptor, respectively. If necessary, kanamycin (50 μg*ml^-1^) was added as selection marker. Arabinose was used in a concentration of 1 mM to express genes cloned in a pBAD vector backbone, 1 mM IPTG was used to express the codon modified *skp* and *surA* versions in the conditional chaperone null mutant.

**Table 3.** Strains, used in this study.

| Strain | Relevant genotype | Reference |
| --- | --- | --- |
| ***S. oneidensis* strains** |  |  |
| *S. oneidensis* MR-1 | WT | 20 |
| *S. oneidensis* MR-1 $\Delta mtrA$ | $\Delta mtrA$ | 51 |
| *S. oneidensis* MR-1 $\Delta mtrA \Delta degP$ | $\Delta mtrA \Delta degP$ | This work |
| *S. oneidensis* MR-1 $\Delta chaperone$ | $\Delta degP::P_{tac}\text{-}skp_{cm}\text{-}surA_{cm}$<br>$\Delta surA skp^*$ | This work |
| ***E. coli* strains** |  |  |
| *E. coli* K-12 pBAD_*mtrB* | pBAD_*mtrB* | This work |
| *E. coli* K-12 pBAD_*mtrAB* | pBAD_*mtrAB* | This work |
| *E. coli* K-12 $\Delta surA$ | $\Delta surA$ | ECGRC (USA) |
| *E. coli* K-12 $\Delta surA$ pBAD_*mtrB* | $\Delta surA mtrB$ | This work |
| *E. coli* K-12 $\Delta skp$ | $\Delta skp$ | ECGRC (USA) |
| *E. coli* K-12 $\Delta skp$ pBAD_*mtrB* | $\Delta skp mtrB$ | This work |
| *E. coli* K-12 $\Delta degP$ | $\Delta degP$ | ECGRC (USA) |
| *E. coli* K-12 $\Delta degP$ pBAD_*mtrB* | $\Delta degP mtrB$ | This work |
| *E. coli* K-12 $\Delta degP \Delta skp$ | $\Delta degP \Delta skp$ | This work |
| *E. coli* K-12 $\Delta degP \Delta skp$ pBAD_*mtrB* | $\Delta degP \Delta skp mtrB$ | This work |
| *E. coli* K-12 $\Delta degP \Delta skp$<br>pBAD_*mtrAB* | $\Delta degP \Delta skp mtrAB$ | This work |

### Cell fractionation

Cells were resuspended in buffer (50 mM HEPES, 150 mM NaCl, pH 7.5), disrupted via french press (1260 psi) or ultrasonication and centrifuged at 6,000 g for 15 minutes. The supernatant was centrifuged at 208,000 g for 1 h to pellet membrane fractions. The latter were homogenized in the same buffer with additional 2% Triton X-100 using a Potter-Elvehjem homogenizer. Soluble and membrane fractions were snap-freezed in 20% glycerol with liquid nitrogen and stored at −80 °C until further use.

### Western blot analysis

Protein concentrations were determined using Roti-Quant (Roth, Germany) with bovine serum albumin as standard. Membrane fractions were loaded on a 12% Mini-PROTEAN TGX protein gel (Bio-Rad, Germany) and separated via SDS-PAGE. Proteins were blotted onto nitrocellulose membranes using Trans-Blot Turbo transfer system with mixed Mw protocol (1,3 A, 25 V for 12 min; Bio-Rad, Germany,). Immunodetection was carried out using primary antibodies raised in rabbit against a peptide specific for MtrB (Gly^23^-Cys^44,35^) or Omp35 (Ser^198^-Tyr^211^) followed by incubation with alkaline phosphatase coupled anti-rabbit antibodies and subsequent detection via the AP detection kit (Bio-Rad, Germany). Imaging was carried out using the ChemiDoc Imaging System and Image Lab Software (Bio-Rad, Germany).

### ß-barrel protein prediction

The complete set of proteins identified within the proteome of the membrane fractions was used as a query for the transmembrane protein topology prediction tool DeepTMHMM^39^. All proteins that were predicted to be monomeric ß-barrel proteins were used for further analysis of the proteome.

### Illumina sequencing

Isolation of genomic DNA, Illumina whole genome sequencing and variant analysis to detect SNPs and InDels was carried out by Eurofins Genomics (Luxembourg) using the Variant analysis pipeline v2.9.15. The raw reads were analyzed with CLC genomics workbench to confirm integrity of the inducible chaperones and the stop codon in *skp*.

*Quantitative Proteomics:*

### Sample Preparation and Digestion

Protein samples were dissolved in 100 mM triethylammonium bicarbonate (TEAB) with 1% (w/v) sodium deoxycholate (SDC), heated at 95 °C for 5 minutes, and then sonicated. Protein concentrations were measured using the Pierce BCA Protein Assay Kit (Thermo Fisher), and samples were normalized to 10 µg protein. Digestion was carried out in 96-well LoBind plates (Eppendorf) using a semi-automated Andrew+ Pipetting Robot (Waters). Proteins were reduced with 10 mM dithiothreitol at 56 °C for 30 minutes (800 rpm) and alkylated with 20 mM iodoacetamide at 37 °C for 30 minutes. Proteins were then immobilized on carboxylate-modified magnetic Sera-Mag SpeedBeads (Cytiva) in a 10:1 bead-to-protein ratio using a 50% acetonitrile (ACN) binding step. Beads were washed twice with 80% ethanol and 100% ACN. Trypsin digestion (Promega; 1:100 enzyme:protein ratio) was performed in 100 mM ammonium bicarbonate at 37 °C overnight (500 rpm). Trypsin was inactivated with 1% trifluoroacetic acid (TFA), and peptide-containing supernatants were collected in new 96-well plates.

### LC-MS/MS Analysis

Peptides were separated via UHPLC (Vanquish Neo, Thermo Fisher) using a two-buffer system (Buffer A: 0.1% FA in water; Buffer B: 0.1% FA in ACN). A trap cartridge (PepMap™ Neo trap cartridge, 5 µm C18, 300 µm × 5 mm, Thermo Fisher) enabled online desalting, followed by separation on a µPAC™ Neo analytical column (C18, 50 cm; Thermo Fisher Scientific). Peptides were eluted over 80 minutes with a gradient from 2% to 30% ACN over 60 minutes.

Mass spectrometry was performed using an Exploris 480 Orbitrap (Thermo Fisher) in data-dependent acquisition (DDA) mode with nano-ESI (1,800 V). MS1 scans covered m/z 350–1400 at 60,000 resolution with a max injection time of 25 ms or 1 × 10⁶ AGC target. The top 20 precursors (intensity > 8 × 10³, charge +2 to +6) were isolated (2 m/z window) and fragmented using HCD (30% NCE). MS2 scans began at m/z 120 with a resolution of 15,000 and AGC target of 1 × 10⁵ or 50 ms injection time. Fragmented precursors were excluded for 30 seconds.

### Data Processing

Raw data were analyzed with Sequest in Proteome Discoverer v3.1 (Thermo Fisher), searching against a reviewed+trEMBL *Shewanella oneidensis* MR-1 database. Carbamidomethylation (C) was set as a fixed modification; oxidation (M), N-terminal pyro-glutamate (Q), and protein N-terminal acetylation were variable. Two missed cleavages were allowed. Only peptides 6–144 amino acids long and meeting a 1% FDR threshold were considered. Quantification was done using the Minora algorithm. Protein abundances were log₂-transformed and normalized to the column median. Proteins identified by match-between-runs only or with fewer than two unique peptides were excluded. The final dataset included proteins consistently detected across all samples.

## Data availability

The mass spectrometry proteomics data have been deposited to the ProteomeXchange Consortium via the PRIDE^53^ partner repository with the dataset identifier PXD064063 and will be made available upon publication.

## Supporting information

Supplemental Dataset S1 - Excel template used to modify codon usage

## Acknowledgments and funding sources

This study was supported by grants from the Deutsche Forschungsgemeinschaft (DFG) (priority program SPP2240 project number 445800740, INST 337/15-1, INST 337/16-1, INST 152/837-1 and INST 152/947-1 FUGG).

We would like to thank Jonna Ram for her assistance with membrane fractionation for quantitative proteomics and Western blot analysis.

Figures 1 – 4 were created with BioRender.com

## Notes

### Competing Interest Statement

The authors have declared no competing interest.

## References

1. Künkele, K. P. et al. The preprotein translocation channel of the outer membrane of mitochondria. Cell 93, 1009–1019 (1998).

2. Baker, K. P., Schaniel, A., Vestweber, D. & Schatz, G. A yeast mitochondrial outer membrane protein essential for protein import and cell viability. Nature 348, 605–609 (1990).

3. Shimizu, S., Narita, M. & Tsujimoto, Y. Bcl-2 family proteins regulate the release of apoptogenic cytochrome *c* by the mitochondrial channel VDAC. Nature 399, 483–487 (1999).

4. Silhavy, T. J., Kahne, D. & Walker, S. The bacterial cell envelope. Cold Spring Harb Perspect Biol 2, a000414 (2010).

5. Sklar, J. G., Wu, T., Kahne, D. & Silhavy, T. J. Defining the roles of the periplasmic chaperones SurA, Skp, and DegP in *Escherichia coli*. Genes Dev 21, 2473–2484 (2007).

6. Chum, A. P., Shoemaker, S. R., Fleming, P. J. & Fleming, K. G. Plasticity and transient binding are key ingredients of the periplasmic chaperone network. Protein Science 28, 1340–1349 (2019).

7. Volokhina, E. B. et al. Role of the Periplasmic Chaperones Skp, SurA, and DegQ in Outer Membrane Protein Biogenesis in *Neisseria meningitidis*. J Bacteriol 193, 1612–1621 (2011).

8. Shen, Q. T. et al. Bowl-shaped oligomeric structures on membranes as DegP’s new functional forms in protein quality control. Proc Natl Acad Sci U S A 106, 4858–4863 (2009).

9. Hennecke, G., Nolte, J., Volkmer-Engert, R., Schneider-Mergener, J. & Behrens, S. The periplasmic chaperone SurA exploits two features characteristic of integral outer membrane proteins for selective substrate recognition. J Biol Chem 280, 23540–23548 (2005).

10. Walton, T. A., Sandoval, C. M., Fowler, C. A., Pardi, A. & Sousa, M. C. The cavity-chaperone Skp protects its substrate from aggregation but allows independent folding of substrate domains. Proc Natl Acad Sci U S A 106, 1772–1777 (2009).

11. Strauch, K. L., Johnson, K. & Beckwith, J. Characterization of *degP*, a gene required for proteolysis in the cell envelope and essential for growth of *Escherichia coli* at high temperature. J Bacteriol 171, 2689–2696 (1989).

12. Chamachi, N. et al. Chaperones Skp and SurA dynamically expand unfolded OmpX and synergistically disassemble oligomeric aggregates. Proc Natl Acad Sci U S A 119, e2118919119 (2022).

13. Narita, S. I., Masui, C., Suzuki, T., Dohmae, N. & Akiyama, Y. Protease homolog BepA (YfgC) promotes assembly and degradation of β-barrel membrane proteins in *Escherichia coli*. Proc Natl Acad Sci U S A 110, E3612–E3621 (2013).

14. Soltes, G. R., Martin, N. R., Park, E., Sutterlin, H. A. & Silhavy, T. J. Distinctive roles for periplasmic proteases in the maintenance of essential outer membrane protein assembly. J Bacteriol 199, e00418–17 (2017).

15. Beliaev, A. S. & Saffarini, D. A. *Shewanella putrefaciens mtrB* encodes an outer membrane protein required for Fe(III) and Mn(IV) reduction. J Bacteriol 180, 6292–6297 (1998).

16. Myers, C. R. & Myers, J. M. MtrB Is Required for Proper Incorporation of the Cytochromes OmcA and OmcB into the Outer Membrane of *Shewanella putrefaciens* MR-1. Appl Environ Microbiol 68, 5585–5594 (2002).

17. Coursolle, D. & Gralnick, J. A. Reconstruction of Extracellular Respiratory Pathways for Iron(III) Reduction in *Shewanella oneidensis* Strain MR-1. Front Microbiol 3, doi:10.3389/fmicb.2012.00056 (2012).

18. Morgan, J. W., Anders, E., Chemical composition of Earth, Venus, and Mercury. Proc Natl Acad Sci U S A 77, 6973–6977 (1980).

19. Richter, K., Schicklberger, M. & Gescher, J. Dissimilatory reduction of extracellular electron acceptors in anaerobic respiration. Appl Environ Microbiol 78, 913–921 (2012).

20. Myers, C. R. & Nealson, K. H. Bacterial manganese reduction and growth with manganese oxide as the sole electron acceptor. Science 240, 1319–1321 (1988).

21. Shi, L., Squier, T. C., Zachara, J. M. & Fredrickson, J. K. Respiration of metal (hydr)oxides by *Shewanella* and *Geobacter*: A key role for multihaem *c*-type cytochromes. Mol Microbiol 65, 12– 20 (2007).

22. Beblawy, S. et al. Extracellular reduction of solid electron acceptors by *Shewanella oneidensis*. Mol Microbiol 109, 571–583 (2018).

23. Liu, J. et al. Identification and characterization of MtoA: A decaheme *c*-type cytochrome of the neutrophilic Fe(ll)-oxidizing bacterium *Sideroxydans lithotrophicus* ES-1. Front Microbiol 3, doi:10.3389/fmicb.2012.00037 (2012).

24. Shi, L., Rosso, K. M., Zachara, J. M. & Fredrickson, J. K. Mtr extracellular electron-transfer pathways in Fe(III)-reducing or Fe(II)-oxidizing bacteria: A genomic perspective. Biochem Soc Trans 40, 1261–1267 (2012).

25. Shi, L. et al. Molecular underpinnings of Fe(III) oxide reduction by *Shewanella oneidensis* MR-1. Front Microbiol 3, doi:10.3389/fmicb.2012.00050 (2012).

26. Shi, L. et al. Direct involvement of type II secretion system in extracellular translocation of *Shewanella oneidensis* outer membrane cytochromes MtrC and OmcA. J Bacteriol 190, 5512– 5516 (2008).

27. Thöny-Meyer, L. Biogenesis of respiratory cytochromes in bacteria. Microbiol Mol Biol Rev 61, 337–376 (1997).

28. Thöny-Meyer, L., Ritz, D. & Hennecke, H. Cytochrome *c* biogenesis in bacteria: a possible pathway begins to emerge. Mol Microbiol 12, 1–9 (1994).

29. Bouhenni, R., Gehrke, A. & Saffarini, D. Identification of genes involved in cytochrome *c* biogenesis in *Shewanella oneidensis*, using a modified *mariner* transposon. Appl Environ Microbiol 71, 4935–4937 (2005).

30. DiChristina, T. J., Moore, C. M. & Haller, C. A. Dissimilatory Fe(III) and Mn(IV) Reduction by *Shewanella putrefaciens* Requires *ferE*, a Homolog of the *pulE* (*gspE*) Type II Protein Secretion Gene. J Bacteriol 184, 142–151 (2002).

31. Hartshorne, R. S. et al. Characterization of an electron conduit between bacteria and the extracellular environment. Proc Natl Acad Sci U S A 106, 22169–22174 (2009).

32. Schicklberger, M., Bücking, C., Schuetz, B., Heide, H. & Gescher, J. Involvement of the *Shewanella oneidensis* Decaheme Cytochrome MtrA in the Periplasmic Stability of the beta-Barrel Protein MtrB. Appl Environ Microbiol 77, 1520–1523 (2011).

33. Philipp, L.-A. et al. Identification of factors limiting the efficiency of transplanting extracellular electron transfer chains in *Escherichia coli*. Appl Environ Microbiol, 0:e00685–25, doi:10.1128/aem.00685-25 (2025).

34. Edwards, M. J., White, G. F., Butt, J. N., Richardson, D. J. & Clarke, T. A. The Crystal Structure of a Biological Insulated Transmembrane Molecular Wire. Cell 181, 665–673 (2020).

35. White, G. F. et al. Rapid electron exchange between surface-exposed bacterial cytochromes and Fe(III) minerals. Proc Natl Acad Sci U S A 110, 6346–6351 (2013).

36. Cao, Y. et al. A synthetic plasmid toolkit for *Shewanella oneidensis* MR-1. Front Microbiol 10, doi:10.3389/fmicb.2019.00410 (2019).

37. Chen, Y. et al. Development of Whole Genome-Scale Base Editing Toolbox to Promote Efficiency of Extracellular Electron Transfer in *Shewanella oneidensis* MR-1. Adv Biol 6, e2101296 (2022).

38. Huang, Q., Szklarczyk, D., Wang, M., Simonovic, M. & von Mering, C. PaxDb 5.0: Curated Protein Quantification Data Suggests Adaptive Proteome Changes in Yeasts. Mol Cell Proteomics 22, doi:10.1016/j.mcpro.2023.100640 (2023).

39. Hallgren, J. et al. DeepTMHMM predicts alpha and beta transmembrane proteins using deep neural networks. bioRxiv [Preprint] doi:10.1101/2022.04.08.487609 (2022).

40. Koronakis, V., Sharff, A., Koronakis, E., Luisi, B. & Hughes, C. Crystal structure of the bacterial membrane protein TolC central to multidrug efflux and protein export. Nature 405, 914–919 (2000).

41. Werner, J., Augustus, A. M. & Misra, R. Assembly of TolC, a Structurally Unique and Multifunctional Outer Membrane Protein of *Escherichia coli* K-12. J Bacteriol 185, 6540–6547 (2003).

42. Theunissen, S. et al. The agglutination protein AggA from *Shewanella oneidensis* MR-1 is a TolC-like protein and forms active channels *in vitro*. Biochem Biophys Res Commun 386, 380– 385 (2009).

43. Chng, S. S., Ruiz, N., Chimalakonda, G., Silhavy, T. J. & Kahne, D. Characterization of the two-protein complex in *Escherichia coli* responsible for lipopolysaccharide assembly at the outer membrane. Proc Natl Acad Sci U S A 107, 5363–5368 (2010).

44. Chimalakonda, G. et al. Lipoprotein LptE is required for the assembly of LptD by the β-barrel assembly machine in the outer membrane of *Escherichia coli*. Proc Natl Acad Sci U S A 108, 2492–2497 (2011).

45. Ruiz, N., Chng, S. S., Hinikera, A., Kahne, D. & Silhavy, T. J. Nonconsecutive disulfide bond formation in an essential integral outer membrane protein. Proc Natl Acad Sci U S A 107, 12245– 12250 (2010).

46. Vertommen, D., Ruiz, N., Leverrier, P., Silhavy, T. J. & Collet, J. F. Characterization of the role of the *Escherichia coli* periplasmic chaperone SurA using differential proteomics. Proteomics 9, 2432–2443 (2009).

47. Schwalm, J., Mahoney, T. F., Soltes, G. R. & Silhavy, T. J. Role for Skp in LptD Assembly in *Escherichia coli*. J Bacteriol 195, 3734–3742 (2013).

48. Zhang, Z., Schwartz, S., Wagner, L. & Miller, W. A Greedy Algorithm for Aligning DNA Sequences. J Comp Biol 7, 203–214 (2000).

49. Datsenko, K. A. & Wanner, B. L. One-step inactivation of chromosomal genes in *Escherichia coli* K-12 using PCR products. Proc Natl Acad Sci U S A 97, 6640–6645 (2000).

50. Shanks, R. M. Q., Caiazza, N. C., Hinsa, S. M., Toutain, C. M. & O’Toole, G. A. *Saccharomyces cerevisiae*-based molecular tool kit for manipulation of genes from gram-negative bacteria. Appl Environ Microbiol 72, 5027–5036 (2006).

51. Schuetz, B., Schicklberger, M., Kuermann, J., Spormann, A. M. & Gescher, J. Periplasmic Electron Transfer via the *c*-Type Cytochromes MtrA and FccA of *Shewanella oneidensis* MR-1. Appl Environ Microbiol 75, 7789–7796 (2009).

52. Gibson, D. G. et al. Enzymatic assembly of DNA molecules up to several hundred kilobases. Nat Methods 6, 343–345 (2009).

53. Perez-Riverol, Y. et al. The PRIDE database at 20 years: 2025 update. Nucleic Acids Res 53, D543–D553 (2025).

54. Bitto, E. & McKay, D. B. Crystallographic structure of SurA, a molecular chaperone that facilitates folding of outer membrane porins. Structure 10, 1489–1498 (2002).

55. Walton, T. A. & Sousa, M. C. Crystal Structure of Skp, a Prefoldin-like Chaperone that Protects Soluble and Membrane Proteins from Aggregation. Mol Cell 15, 367–374 (2004).

56. Krojer, T., Garrido-Franco, M., Huber, R., Ehrmann, M. & Clausen, T. Crystal structure of DegP (HtrA) reveals a new protease-chaperone machine. Nature 416, 455–459 (2002).

