## Supplemental Information for "Identification of chaperone-independent outer membrane proteins and MtrA-assisted MtrB folding in *Shewanella oneidensis*"

**This PDF file includes:**

Figures S1 to S2
Tables S1 to S2

**Other supporting materials for this manuscript include the following:**

Dataset S1

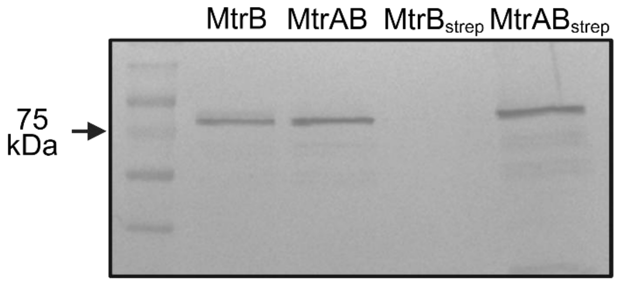


*Figure S1* Western blot analysis of *E. coli* membrane fractions using an antibody specific for MtrB after expression of either MtrB or a strep-tagged MtrB version with and without co-expression of MtrA. Only the strep-tagged MtrB is degraded in the absence of MtrA.

ATTGCCTGGCAAGGTATCAAAAAAGGTGTCCATCAGATAGGTTTTGCCGCGGCCAACGCCCCCC
CAAAGATAAAGTCCTTTTGTAGGCGCGGGCGCAGATGTTAACCCAAAAGAGGTAAATAATTTACC
AAACAAAGAGTTAGGCTCTTCGGCTGCGGTTAAATCCTCATAAACCCGTTGCAGCGCCTTAACAG
CAATTTCCTGCGCCGGATCATGGGAAAAACCATCTCGAGTAAGATCTTTTTGGTAATGCTGCCAA
GGGCTTAACTGGGGCACGTTATAATTCTCTTATATCCTTTATTGAAACTGCGTCGATATTAGCACG
GCCTAGATGACACAGCATATATTTATACCCAAGCGACCTGAATTCAGCATTTTTAAATCACCTAGT
AATAAGTACTGCAACACCACTAAAACCATGTATAAAATGATTAAAACTGATTTATAAAGGACTATGC
TGTTAGCTAATTAAACTGAACGGTTTGGATCATTTAACACTTGACACCATCGAATGGTGCAAAACC
TTTCGCGGTATGGCATGATAGCGCCCGGAAGAGAGTCAATTCAGGGTGGTGAATGTGAAACCAG
TAACGTTATACGATGTCGCAGAGTATGCCGGTGTCTCTTATCAGACCGTTTCCCGCGTGGTGAAC
CAGGCCAGCCACGTTTCTGCGAAAACGCGGGAAAAAGTGGAAGCGGCGATGGCGGAGCTGAAT
TACATTCCCAACCGCGTGGCACAACAACTGGCGGGCAAACAGTCGTTGCTGATTGGCGTTGCCA
CCTCCAGTCTGGCCCTGCACGCGCCGTCGCAAATTGTCGCGGCGATTAAATCTCGCGCCGATCA
ACTGGGTGCCAGCGTGGTGGTGTCGATGGTAGAACGAAGCGGCGTCGAAGCCTGTAAAGCGGC
GGTGCACAATCTTCTCGCGCAACGCGTCAGTGGGCTGATCATTAACTATCCGCTGGATGACCAG
GATGCCATTGCTGTGGAAGCTGCCTGCACTAATGTTCCGGCGTTATTTCTTGATGTCTCTGACCA
GACACCCATCAACAGTATTATTTTCTCCCATGAAGACGGTACGCGACTGGGCGTGGAGCATCTGG
TCGCATTGGGTCACCAGCAAATCGCGCTGTTAGCGGGCCCATTAAGTTCTGTCTCGGCGCGTCT
GCGTCTGGCTGGCTGGCATAAATATCTCACTCGCAATCAAATTCAGCCGATAGCGGAACGGGAA
GGCGACTGGAGTGCCATGTCCGGTTTTCAACAAACCATGCAAATGCTGAATGAGGGCATCGTTCC
CACTGCGATGCTGGTTGCCAACGATCAGATGGCGCTGGGCGCAATGCGCGCCATTACCGAGTCC
GGGCTGCGCGTTGGTGCGGATATCTCGGTAGTGGGATACGACGATACCGAAGACAGCTCATGTT
ATATCCCGCCGTTAACCACCATCAAACAGGATTTTCGCCTGCTGGGGCAAACCAGCGTGGACCG
CTTGCTGCAACTCTCTCAGGGCCAGGCGGTGAAGGGCAATCAGCTGTTGCCCGTCTCACTGGTG
AAAAGAAAAACCACCCTGGCGCCCAATACGCAAACCGCCTCTCCCCGCGCGTTGGCCGATTCAT
TAATGCAGCTGGCACGACAGGTTTCCCGACTGGAAAGCGGGCAGTGAGCGCAACGCAATTAATG
TAAGTTAGCTCACTCATTAGGCACAATTCTCATGTTTGACAGCTTATCATCGACTGCACGGTGCAC
CAATGCTTCTGGCGTCAGGCAGCCATCGGAAGCTGTGGTATGGCTGTGCAGGTCGTAAATCACT
GCATAATTCGTGTCGCTCAAGGCGCACTCCCGTTCTGGATAATGTTTTTTGCGCCGACATCATAA
CGGTTCTGGCGAAATATTCTGAAATGAGCTGTTGACAATTAATCATCGGCTCGTATAATGTGTGGA
ATTGTGAGCGGATAACAATTAATAATTTTGTTTAACTTTAAGAAGGAGATATAATGGTCAACCGCG
CATTAGTTACTTTGGCGCTGTTAGGCGCCCCACTGGCTGCTCAGGCCGAGAACATCGCGGTAGT
TGACATGGGTGCGGTTTTTGAACAATTACCTCAACGTGAACAGATTATGCAATCTTTAAAATCTGA
ATTTGGCGATCGTATGAGCGAGGTACAAAAAATGCAGGAAGAAATGCGTTCTTTAATGGAAAAAC
AACAACGCGACGGTGCTCTGATGAACGATACTCAGAAAACGGAATTAGTGCGTAAAATGGAGGCT
CTGAAATCTGAGTACCAATTAAAGGGCAAAGCTTTAGATGAAGATTTACGTCGCCGTCAAGGCGA
GGAACAGAATAAGTTATTAGTTAAAGTTCAAAAAGCAATCAATACGATTGCGGAAAAGGAGAAGTA
TGACCTGGTGTTACAACGTGGTGCCGTGATCTATGTTAAACCTAACGCGGATATTTCAGGTAAAG
TTGTGGAAGCATTGAGTAAGGGTAAATAACACTCTGCGCATCACGACACTGTTTTATGAACAGCA
CTAAATAAAAGGAGTGAAGGGAAATATGAAGCCATCTAAGCACTTAATCTTCGCACTCTTCGCTCT
GGCAATTTCACAACCCACCATGGCGGCTCCTCAACCCCTGGATCGCGTCGCTGTCCAGATTAATG
ATGGCATCGTTCTGGAGTCTGAAATTACTAACATGATCGATACAGTGAAAGCGAACGCAAAAGCG
GCGAACCAATCCTTACCATCGGATTCTGCCTTACGTACCCAAGTCATTGAAAGACTCATCTTGACC
CGCTTACAGCTCCAAATGGCCGACCGTATCGGCTTACACATCGGCGACCTGCAACTCGATCAGG
CCATCGAGAATATCGCCCGCGAACAAAAAATGACCGTGGCCCAGATGCAACAAAAGATCGCTTCA
GAAGGGATATCTTTTTCACAATACCGTGAGCAGCTTCGCGAAGAGATTACTCTCGGCGAAATCCA
ACGCATCCAGGTACAACGTCGCATTCAGGTCAGCCCTCAAGAAATTACCGGCCTAGTTAAATTAA
TTCAAGAGCAAGGCATGAAAGATGTCGAATATCAGATTGGCCACATTTTAATCGATGTCCCGAACA
ACCCAACTTCTGAACAACTGGAAGCGAGCTCAAAACGTGCCAACGCGGTTCTAGAGCGCCTCAA
ATCTGGCGAAGACTTTCGTCGCACCGCGATTGCCAGTAGCAGTGGCCCAAAGGCTCTAGAAGGC
GGCATTTGGGACTATATGAATATTAATGAAATGCCCACTTTATTCGCGGAGGTGATCAATGGCGC
CAAAAAAGGCGACATCATTGGCCCCATCAAAACCGGCGCTGGCTTTCACATTATTAAAATTATGGA
CGCACGTGGCCTCCAGACTAAGGAAATCGAAGAAGTGCGAGCACGCCATATCCTTCTGAAACCC
AGCCCAATCTTGAGTGAAGACCGTGCAAAAGCTATGCTGGAACAATTTCTAAAACAAATCCGTAGT
GGCGAAGCAAAATTCGAGGACCTCGCGCGCCAATACAGTGAAGACCCCGGCAGTGCCACCAAA
GGCGGCGAACTTGGCTGGGCTGAACCTTCTATCTATGTTCCCGAATTCGCCCAGACACTGAACTC
TCTCTCACCCGACCAAATTTCTGAACCATTTCGCACAACCCATGGCTGGCATATTACCCAACTCGA
GGAACGTAGGAAGACTGACGCAACGGACCAATTCAATACCAATCGCGCCCATCAATTAATCTTTC
GCCGTAAGTTCAACGAAGAGTTACAGAATTGGCTTGATGAAATGCGCGCGGATGCTTATATCGAA
GTGTTCCAACCTGAAAGTAACAGAGGCTAAAACATGTTAATCTAACTCACCTTTTTCAATATTTAGC
CTGCCATGACAATAAAAGACACCCTGTTATATCTCGGTAAAGCCGTGCTTTTCGGCCTTATTATGG
CAGCCATGTTTTTGTTAGTAACTCATTACTTCGACAATAAGAGTCTTGGCAACTCATTACTACAAAA

CCGCGGTAACAACACGGTTGAACTCTCCTTTGCCAAGGCCGTGCGTCGCGCAGCCCCTGCCGTC
GTCAATATTTACAGCTTGAGTATCGATCAGAGCCGTCCATTGAATTCTGGCTCACTCCAAGGTCTA
GGATCTGGGGTGATCATGAGTAAAGAAGGTTATATTCTCACTAATTATCATGTGATTAAAAAAGCC
GATGAGATCGTCGTCGCGCTGCAAGATGGCCGCAAGTTCACCTCTGAAGTGGTGGGGTTTGATC
CCGAAACCGATCTTTCAGTGCTTAAGATCGAAGGCGATAATCTCCCCACTGTGCCAGTCAATCTC
GACAG

*Figure S2* DNA sequence used for the integration of codon modified versions of *skp* and *surA* in the *degP* locus of *S. oneidensis*. The sequence contains 500 bp upstream of *degP* (teal), the repressor *lacI* (red), inducible promoter pTac (green), codon modified version of *skp* (pink) and *surA* (yellow) as well as 500 bp downstream of *degP* (teal).

*Table S1* Frequency of codons used in the modified and native versions of *surA* and *skp*.

| | Frequency of Codons | | | |
|---|---|---|---|---|
| **Codon** | ***surA* modified** | ***surA* native** | ***skp* modified** | ***skp* native** |
| **AAA** | 0.04 | 0.04 | 0.07 | 0.07 |
| **AAC** | 0.02 | 0.02 | 0.02 | 0.02 |
| **AAG** | 0.02 | 0.02 | 0.03 | 0.03 |
| **AAT** | 0.02 | 0.02 | 0.01 | 0.01 |
| **ACA** | 0.01 | 0.01 | 0.00 | 0.00 |
| **ACC** | 0.03 | 0.03 | 0.00 | 0.00 |
| **ACG** | 0.00 | 0.00 | 0.01 | 0.01 |
| **ACT** | 0.01 | 0.01 | 0.01 | 0.01 |
| **AGA** | 0.00 | 0.00 | 0.00 | 0.00 |
| **AGC** | 0.01 | 0.01 | 0.01 | 0.01 |
| **AGG** | 0.00 | 0.00 | 0.00 | 0.00 |
| **AGT** | 0.02 | 0.02 | 0.01 | 0.01 |
| **ATA** | 0.00 | 0.00 | 0.00 | 0.00 |
| **ATC** | 0.05 | 0.04 | 0.02 | 0.02 |
| **ATG** | 0.03 | 0.03 | 0.05 | 0.05 |
| **ATT** | 0.04 | 0.05 | 0.02 | 0.02 |
| **CAA** | 0.06 | 0.06 | 0.06 | 0.06 |
| **CAC** | 0.01 | 0.01 | 0.00 | 0.00 |
| **CAG** | 0.03 | 0.02 | 0.03 | 0.03 |
| **CAT** | 0.01 | 0.01 | 0.00 | 0.00 |
| **CCA** | 0.01 | 0.02 | 0.01 | 0.01 |
| **CCC** | 0.02 | 0.01 | 0.00 | 0.00 |
| **CCG** | 0.00 | 0.00 | 0.00 | 0.00 |
| **CCT** | 0.01 | 0.01 | 0.01 | 0.01 |
| **CGA** | 0.00 | 0.00 | 0.00 | 0.00 |
| **CGC** | 0.03 | 0.03 | 0.02 | 0.02 |
| **CGG** | 0.00 | 0.00 | 0.00 | 0.00 |
| **CGT** | 0.03 | 0.03 | 0.04 | 0.04 |
| **CTA** | 0.01 | 0.01 | 0.00 | 0.00 |
| **CTC** | 0.02 | 0.02 | 0.00 | 0.00 |
| **CTG** | 0.02 | 0.02 | 0.03 | 0.03 |
| **CTT** | 0.01 | 0.01 | 0.00 | 0.00 |
| **GAA** | 0.07 | 0.07 | 0.07 | 0.07 |
| **GAC** | 0.03 | 0.02 | 0.02 | 0.02 |
| **GAG** | 0.02 | 0.02 | 0.04 | 0.04 |
| **GAT** | 0.02 | 0.03 | 0.03 | 0.03 |
| **GCA** | 0.02 | 0.02 | 0.02 | 0.02 |
| **GCC** | 0.03 | 0.02 | 0.02 | 0.02 |
| **GCG** | 0.02 | 0.03 | 0.03 | 0.03 |
| **GCT** | 0.02 | 0.02 | 0.03 | 0.03 |
| **GGA** | 0.00 | 0.00 | 0.00 | 0.00 |

| | | | | |
|---|---|---|---|---|
| **GGC** | 0.06 | 0.02 | 0.02 | 0.02 |
| **GGG** | 0.00 | 0.00 | 0.00 | 0.00 |
| **GGT** | 0.00 | 0.03 | 0.03 | 0.03 |
| **GTA** | 0.00 | 0.00 | 0.01 | 0.01 |
| **GTC** | 0.01 | 0.01 | 0.01 | 0.01 |
| **GTG** | 0.01 | 0.01 | 0.02 | 0.02 |
| **GTT** | 0.01 | 0.01 | 0.04 | 0.04 |
| **TAA** | 0.00 | 0.00 | 0.00 | 0.00 |
| **TAC** | 0.00 | 0.00 | 0.01 | 0.01 |
| **TAG** | 0.00 | 0.00 | 0.00 | 0.00 |
| **TAT** | 0.01 | 0.01 | 0.01 | 0.01 |
| **TCA** | 0.01 | 0.01 | 0.01 | 0.01 |
| **TCC** | 0.00 | 0.00 | 0.00 | 0.00 |
| **TCG** | 0.00 | 0.00 | 0.00 | 0.00 |
| **TCT** | 0.02 | 0.02 | 0.02 | 0.02 |
| **TGA** | 0.00 | 0.00 | 0.00 | 0.00 |
| **TGC** | 0.00 | 0.00 | 0.00 | 0.00 |
| **TGG** | 0.01 | 0.01 | 0.00 | 0.00 |
| **TGT** | 0.00 | 0.00 | 0.00 | 0.00 |
| **TTA** | 0.02 | 0.02 | 0.07 | 0.07 |
| **TTC** | 0.02 | 0.01 | 0.00 | 0.00 |
| **TTG** | 0.00 | 0.00 | 0.01 | 0.01 |
| **TTT** | 0.01 | 0.02 | 0.01 | 0.01 |

*Table S2* Mean normalized $log_2$ abundances and $log_2$ fold changes of the reliable quantifiable ß-barrel proteins after pre-culture and samples t1 and t2. Mean abundances are calculated from three independent replicates with standard deviation given in brackets. The $log_2$ fold changes, with p-values given in brackets, represent the difference between mean $log_2$ abundances of not induced versus induced cells.

| **Gene name** | **pre-culture** | **t1** | | | **t2** | | |
|---|---|---|---|---|---|---|---|
| | $log_2$ abundance induced cells | Mean $log_2$ abundance - not induced cells | Mean $log_2$ abundance - induced cells | $log_2$ fold change (p-value) | Mean $log_2$ abundance - not induced cells | Mean $log_2$ abundance - induced cells | $log_2$ fold change (p-value) |
| ***lptD*** | 1.66 | 1.65 (0.26) | 3.17 (0.31) | -1.52 (0.0058) | 1.24 (0.64) | 2.24 (0.14) | -1.00 (0.0986) |
| ***mtrB*** | 4.57 | 5.09 (0.19) | 6.06 (0.09) | -0.97 (0.0028) | 5.14 (0.04) | 5.51 (0.75) | -0.37 (0.5300) |
| ***dmsF*** | 1.68 | 1.62 (0.20) | 4.64 (0.17) | -3.02 (0.0001) | 1.99 (0.11) | 4.02 (0.49) | -2.03 (0.0048) |
| ***wbfB*** | 4.26 | 2.69 (0.08) | 4.67 (0.15) | -1.98 (0.0001) | 2.15 (0.51) | 4.35 (0.06) | -2.20 (0.0038) |
| ***SO_2469*** | 3.98 | 4.59 (0.30) | 8.07 (0.12) | -3.48 (0.0001) | 7.07 (0.16) | 7.73 (0.49) | -2.58 (0.0018) |
| ***SO_2907*** | 6.93 | 4.37 (0.12) | 8.15 (0.16) | -3.78 (0.0000) | 4.64 (0.09) | 7.64 (0.07) | -3.00 (0.0000) |
| ***SO_3514*** | -0.96 | -0.64 (0.17) | 0.94 (0.31) | -1.59 (0.0031) | -0.65 (0.48) | -0.11 (0.24) | -0.54 (0.2316) |
| ***SO_2427*** | 6.72 | 5.20 (0.22) | 7.07 (0.16) | -1.87 (0.0006) | 5.97 (0.09) | 6.78 (0.29) | -0.81 (0.0200) |
| ***SO_0719*** | 4.00 | 0.96 (0.38) | 2.87 (0.15) | -1.91 (0.0028) | 0.41 (0.18) | 1.69 (0.43) | -1.28 (0.0176) |
| ***SO_2715*** | 1.29 | -1.36 (0.55) | 1.34 (0.17) | -2.70 (0.0027) | -1.60 (0.25) | 0.67 (0.32) | -2.26 (0.0014) |
| ***SO_4422*** | 3.84 | -0.56 (0.94) | 1.06 (0.42) | -1.61 (0.0907) | -1.28 (0.26) | 0.73 (0.31) | -2.01 (0.0023) |
| ***btuB*** | 5.68 | 3.89 (0.27) | 6.11 (0.20) | -2.21 (0.0008) | 6.11 (0.20) | 5.49 (0.18) | -1.35 (0.0006) |

| | | | | | | | |
|---|---|---|---|---|---|---|---|
| ***plpD*** | 2.22 | 0.65 (0.13) | 3.37 (0.22) | -2.72 (0.0001) | 3.37 (0.22) | 2.62 (0.13) | -2.89 (0.0249) |
| ***SO_4685*** | 2.48 | 0.69 (0.31) | 2.72 (0.27) | -2.03 (0.0022) | 0.51 (0.53) | 2.62 (0.06) | -2.12 (0.0049) |
| ***bamA*** | 5.65 | 4.90 (0.21) | 6.13 (0.25) | -1.22 (0.0062) | 6.13 (0.25) | 5.53 (0.24) | -0.77 (0.0121) |
| ***SO_2114*** | -1.13 | 0.11 (0.26) | 1.22 (0.30) | -1.12 (0.0168) | -0.75 (1.60) | 0.79 (0.22) | -1.54 (0.2482) |
| ***SO_1821*** | 5.92 | 6.92 (0.37) | 8.07 (0.16) | -1.14 (0.0156) | 1.22 (0.30) | 7.40 (0.20) | -0.40 (0.0590) |
| ***SO_0564*** | 1.59 | 2.21 (0.35) | 4.53 (0.10) | -2.32 (0.0008) | 4.30 (0.09) | 3.58 (0.04) | -1.05 (0.0001) |
| ***SO_3060*** | -2.73 | -3.71 (0.33) | -1.16 (0.12) | -2.55 (0.0005) | 8.07 (0.16) | -1.52 (0.15) | -1.05 (0.0148) |
| ***SO_1412*** | -0.04 | 0.28 (0.18) | 1.93 (0.17) | -1.65 (0.0008) | 0.63 (0.27) | 1.65 (0.34) | -1.53 (0.0056) |
| ***SO_0312*** | 2.25 | 1.21 (0.29) | 4.30 (0.09) | -3.09 (0.0001) | 0.71 (0.19) | 3.08 (0.10) | -2.37 (0.0001) |
| ***SO_3298*** | -2.17 | -1.39 (0.29) | 0.63 (0.27) | -2.02 (0.0019) | -1.53 (0.24) | -0.06 (0.56) | -1.47 (0.0275) |
| ***omp35*** | 10.39 | 9.61 (0.33) | 11.86 (0.16) | -2.25 (0.0009) | 9.81 (0.11) | 11.46 (0.26) | -1.65 (0.0013) |
| ***SO_3099*** | 6.19 | 5.09 (0.20) | 5.88 (0.19) | -0.79 (0.0159) | 4.27 (0.11) | 5.33 (0.65) | -1.06 (0.0871) |
| ***SO_3276*** | -1.72 | -0.44 (0.48) | 1.04 (0.22) | -1.48 (0.0163) | -0.15 (0.09) | 0.17 (0.23) | -0.33 (0.1399) |
| ***SO_2736*** | -0.96 | -1.31 (0.29) | -0.41 (0.41) | -0.91 (0.0626) | -1.50 (0.20) | -0.79 (0.84) | -0.70 (0.3150) |
| ***mipA*** | -1.61 | -1.94 (0.42) | -0.53 (0.44) | -1.41 (0.0309) | -1.86 (0.34) | -0.99 (0.17) | -0.88 (0.0310) |
| ***ompK*** | 6.23 | 5.42 (0.37) | 7.25 (0.25) | -1.83 (0.0045) | 5.17 (0.18) | 6.77 (0.43) | -1.59 (0.0082) |
| ***SO_1045*** | 1.58 | 0.68 (0.37) | 2.44 (0.25) | -1.77 (0.0051) | 0.86 (0.15) | 1.36 (0.09) | -0.50 (0.0171) |
| ***SO_3931*** | -2.76 | -1.00 (0.54) | -0.94 (0.32) | -0.06 (0.8948) | -1.55 (0.50) | -1.65 (0.71) | 0.10 (0.8824) |

| | | | | | | | |
|---|---|---|---|---|---|---|---|
| ***SO_4131*** | 0.13 | -1.47 (0.42) | 1.01 (0.17) | -2.48 (0.0015) | -1.52 (0.07) | -0.30 (0.17) | -1.22 (0.0008) |
| ***SO_0934*** | -1.19 | 0.43 (0.33) | 0.26 (0.31) | 0.17 (0.6210) | -0.43 (0.47) | -0.41 (0.99) | -0.02 (0.9829) |
| ***SO_3905*** | 1.69 | 2.31 (0.47) | 3.26 (0.22) | -0.95 (0.0603) | 1.78 (0.16) | 2.85 (0.70) | -1.06 (0.1046) |
| ***yfaZ***, | 3.57 | 3.66 (0.19) | 3.61 (0.33) | 0.05 (0.8473) | 3.26 (0.22) | 2.86 (0.62) | 0.46 (0.3837) |
| ***SO_3545*** | 5.72 | 6.17 (0.33) | 7.10 (0.27) | -0.93 (0.0359) | 6.46 (0.21) | 6.69 (0.77) | -0.23 (0.7094) |
| ***SO_3810*** | -0.26 | 1.13 (0.24) | -0.27 (0.38) | 1.40 (0.0120) | 0.50 (0.11) | -1.15 (0.77) | 1.65 (0.0398) |

*Dataset S1 (separate file)* Excel template used to modify the codon usage of *skp* and *surA*.
